# Multimodal gradients across mouse cortex

**DOI:** 10.1101/393215

**Authors:** Ben D. Fulcher, John D. Murray, Valerio Zerbi, Xiao-Jing Wang

## Abstract

The primate cerebral cortex displays a hierarchy that extends from primary sensorimotor to association areas, supporting increasingly integrated function underpinned by a gradient of heterogeneity in the brain’s microcircuits. The extent to which these hierarchical gradients are unique to primate or may reflect a conserved mammalian principle of brain organization remains unknown. Here we report the topographic similarity of large-scale gradients in cytoarchitecture, gene expression, interneuron cell densities, and long-range axonal connectivity, which vary from primary sensory through to prefrontal areas of mouse cortex, highlighting an underappreciated spatial dimension of mouse cortical specialization. Using the T1w:T2w magnetic resonance imaging map as a common spatial reference for comparison across species, we report interspecies agreement in a range of large-scale cortical gradients, including a significant correspondence between gene transcriptional maps in mouse cortex with their human orthologs in human cortex, as well as notable interspecies differences. Our results support the view of systematic structural variation across cortical areas as a core organizational principle that may underlie hierarchical specialization in mammalian brains.

---

Across the brain, cortical microcircuits vary in their cytoarchitecture (1–3), myeloarchitecture (3–5), dendritic and synaptic structure (6–13), and the size, density, and laminar distribution of distinct cell types (1, 3, 14–16). Many of these microstructural properties vary across the cortex continuously as spatial gradients (1–3, 17–20) that shape the specialized functional capabilities of different cortical areas (7), through variations in plasticity (5), inhibitory control (21), and electrophysiological properties (6, 10). Prominent gradients follow a hierarchy of increasing functional integration, from primary sensory to transmodal association areas in primate cortex, pointing to their role in shaping functional specialization along the cortical hierarchy (22–25). The mouse cortex is relatively uniform compared to the highly differentiated primate cortex (10, 26–28), but recent evidence has nevertheless pointed to a global hierarchy of mouse cortical areas (29, 30). It remains unknown whether the hierarchical gradients of primate cortex exist and play a similar role in functional specialization in the mouse, and may therefore represent a conserved property of mammalian brain organization.

The mouse is an ideal model to investigate gradients of cortical microstructure, with experimental datasets from diverse modalities available in standardized anatomical reference spaces (31). Cortical maps of a wide range of properties, many of which are unavailable in human, have been measured in mouse, including: (i) gene expression with approximate genome-wide coverage (32), (ii) interneuron cell densities (29), (iii) tract-traced axonal connectivity (30, 33–36), (iv) cytoarchitecture (37), (v) cell/neuron density (38–40), and (vi) resting-state function magnetic resonance imaging (fMRI) (41–43). Existing work has demonstrated an association between pairs of these measurements (42–44), but these data have not previously been characterized together from the viewpoint of macroscopic gradients.

The T1w:T2w ratio is a noninvasive MRI measurement that has been measured in mouse, macaque, and human, providing a common spatial reference map for linking large-scale cortical gradients across species. Commonly interpreted as a marker of intracortical myelin content (45), T1w:T2w is also sensitive to a wide range of other microstructural properties (46). In macaque cortex, T1w:T2w is strongly correlated to the established structural hierarchy of feedforward-feedback interareal laminar projections (47), and in human cortex it follows dominant dominant gene transcriptional gradients, positioning it as a strong candidate marker of hierarchical specialization (25). Here we show for the first time that gradients of diverse properties of mouse cortex exhibit a common spatial patterning along a candidate functional hierarchy. We use T1w:T2w as a common spatial reference to demonstrate a correspondence of gradients of cytoarchitecture between mouse, macaque, and human, and with transcriptional maps of ortholog genes between mouse and human. Our results reveal an interspecies conservation of cortical gradients that may shape the functional specialization of mammalian cortical circuits, consistent with systematic structural variation across cortical areas as a core organizing principle (48).

## Results

We analyzed the spatial maps of diverse cortical properties across 40 areas of the Allen Reference Atlas (ARA) (49), shown in Figs 1A,B. The similarity between two spatial maps was quantified as the Spearman rank correlation coefficient, *ρ*, across as many cortical areas as could be matched between a given pair of modalities (40 unless otherwise specified). Corrected *p*-values, *p*_corr_, are calculated using the method of Benjamini and Hochberg (50).

**Fig. 1.**
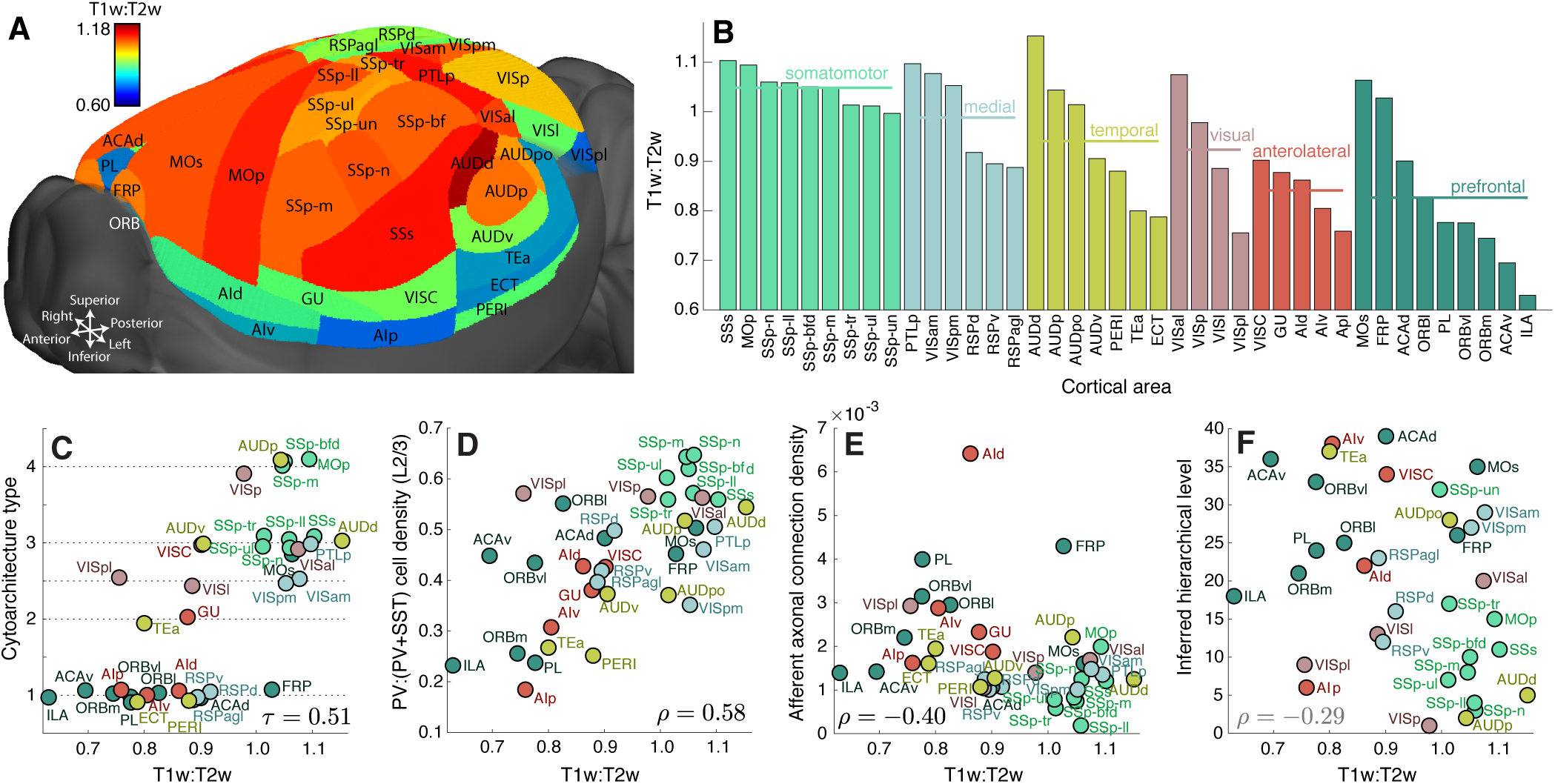
The spatial map of the MRI measurement, T1w:T2w, is correlated with diverse structural properties. **A** Variation in T1w:T2w across mouse cortical areas (visualization through medial sections are in Fig. S3). **B** T1w:T2w broadly decreases across families of connectivity-based groupings of cortical areas (30), from somatomotor areas through to anterolateral and prefrontal areas. Groups are ordered by decreasing T1w:T2w, as are areas within each group. Scatter plots are shown for T1w:T2w (horizontal axis) versus: **C** cytoarchitecture type (37), **D** Relative density of PV:(PV+SST) cells in L2/3 (29), **E** Sum of incoming inter-areal axonal projection weight (33), and **F** Hierarchical level inferred from feedforward–feedback laminar projection patterns (30). Circles are colored according to the connectivity modules in **B** and have sizes scaled by T1w:T2w. A small amount of vertical noise has been added to points in **C** to aid visualization (cytoarchitecture type is discrete).

### Cortical gradients follow T1w:T2w

We first investigated whether the T1w:T2w map is informative of functionally relevant macroscopic gradients of cortical variation in mouse, as it is in macaque and human (25). As shown in Fig. 1A, the T1w:T2w map of the mouse cortex displays non-trivial anatomical specificity on top of a broad spatial embedding: increasing along the inferior–superior axis, |*ρ*| = 0.53 (*p* = 5 × 10^−4^), perhaps in part reflecting a neurodevelopmental gradient of cortical maturation (51) (see Supporting Information for more details). Figure 1B shows that T1w:T2w broadly decreases across five spatially-localized connectivity modules (30), from somatomotor to prefrontal areas, consistent with a decreasing trend from primary unimodal through to transmodal areas in macaque and human (25). T1w:T2w is not significantly correlated to variation in cell density, *ρ* = 0.21 (*p* = 0.2) (39), nor neuron density, *ρ* = −0.08 (*p* = 0.6, across 39 matching cortical areas) (40), making it well positioned to capture density-independent differences in neural architecture (1) (alternative data sources gave consistent results, see Supplementary Information).

Intracortical myelin, which T1w:T2w is sensitive to (45), increases with laminar differentiation (4, 5), with areas lower in the cortical hierarchy exhibiting a greater degree of laminar elaboration (22). Accordingly, T1w:T2w captures the variation in cytoarchitecture in macaque, *τ* = 0.87 (across eight cytoarchitectonic types) and human, *ρ* = 0.74 (using layer IV gene markers) (25). As shown in Fig. 1C, we report a similar trend in mouse cortex, with T1w:T2w increasing from dysgranular to eulaminar areas, Kendall’s *τ* = 0.51 (*p* = 2 × 10^−6^, using five cytoarchitectonic categories assigned to 38 matching cortical areas (37)).

Interneuron densities vary across mouse cortical areas (29). In layer 2/3, sensory-motor areas contain a greater proportion of parvalbumin-containing (PV) interneurons, while association and frontal areas contain a greater proportion of somatostatin-containing (SST) interneurons; the ratio PV : (PV + SST) orders cortical areas along a candidate functional hierarchy (29). We computed the correlation between T1w:T2w and layer 2/3 density of each of three measured interneuron cell types: PV, SST, and vasoactive intestinal peptide-containing (VIP) cells, as well as the proposed hierarchy marker, PV:(PV+SST) (29), across 36 matching cortical areas. PV:(PV+SST) is positively correlated with T1w:T2w, *ρ* = 0.58 (*p*_corr_ = 5 × 10^−4^, correcting for four independent comparisons), as shown in Fig. 1D. The result is consistent with existing studies that have found a covariation of PV neuron density with myelin content (5) and the degree of laminar differentiation (15), leading to a characteristic variation of inhibitory control (2, 7, 48). Given a loose interpretation of SST interneurons as having a putative ‘input-modulating’ function relative to the more ‘output-modulating’ PV interneurons (29), this trend is also consistent with more functionally integrative areas (lower T1w:T2w) requiring greater input control. We also compared the interareal variation of T1w:T2w to cell densities (mm^−3^) reported by Erö et al. (40) for glia, excitatory cells, inhibitory cells, modulatory cells, astrocytes, oligodendrocytes, and microglia across 39 matching cortical areas. We found a significant correlation between T1w:T2w and the density of glia, *ρ* = 0.48 (*p*_corr_ = 0.01); inhibitory cells, *ρ* = −0.47 (*p*_corr_ = 0.01); microglia, *ρ* = 0.37 (*p*_corr_ = 0.04); and oligodendrocytes, *ρ* = 0.37 (*p*_corr_ = 0.04) (scatter plots shown in Fig. S6).

We next investigated whether T1w:T2w is related to properties of interareal axonal connectivity, measured using viral tract tracing (33), focusing on the normalized axonal connection density projected to (weighted in-degree, 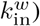 and from (weighted out-degree, 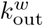) each cortical area. Across 38 matching areas, T1w:T2w is significantly correlated with 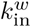, *ρ* = −0.40 (*p*_corr_ = 0.03, correcting for two independent comparisons), plotted in Fig. 1E, but not with 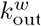, *ρ* = 0.14 (*p*_corr_ = 0.4). This trend reflects a greater aggregate strength of axonal inputs in more functionally integrative areas (with lower T1w:T2w).

Comprehensive data on laminar-specific intracortical projection patterns have recently been used to assign candidate hierarchical levels to mouse cortical areas (30). The mouse cortex does not fit neatly into a global structural hierarchy, with a hierarchy score of just 0.126 (where 0 is non-hierarchical and 1 is perfectly hierarchical) (30). Here we found a weak negative correlation between T1w:T2w and hierarchical level in mouse *ρ* = −0.29 (*p* = 0.09), as shown in Fig. 1F, in contrast to the strong negative correlation reported in macaque, *ρ* = −0.76 (25).

### Gene transcription

Gene transcriptional maps of the mouse brain, measured with cellular resolution using *in situ* hybridization, form the Allen Mouse Brain Atlas (AMBA) (32). We developed and applied stringent quality criteria to obtain cortical expression maps for 4181 genes (listed in Supporting Information). We first focus on a selected set of 86 receptor subunit and cell-type marker genes (54) (see Supporting Information). After correcting for testing multiple independent hypotheses (50) (a conservative correction due to the high intercorrelation between many of these genes), we found that the transcriptional maps of 24 genes display are significantly correlated with T1w:T2w (*p*_corr_ < 0.05; |*ρ*| ≥0.39), ranging from glutamate receptor subunits (*Grin3a, Grin2d, Grik1, Grik2, Grik4, Grm2, Grm5*); serotonin receptor subunits (*Htr1a, Htr2c*, and *Htr5b*); interneuron cell-type markers (*Pvalb* and *Calb2*); the myelin marker, *Mobp*; and a range of other receptor subunit genes: *Trhr, Mc4r, Chrm5, Galr2, Hcrtr2, Hcrtr1, P2ry12, P2ry14, Cnr1, Oxtr, P2ry2* (see Table S2 for full results). Correlation coefficients between T1w:T2w and transcriptional levels of a selected subset of glutamate receptor subunit and interneuron marker genes are plotted in Fig. 2A. The strong negative correlation between T1w:T2w and *Grin3a* expression, *ρ* = −0.63 (*p*_corr_ = 5 × 10^−4^), is shown in Fig. 2B.

**Fig. 2.**
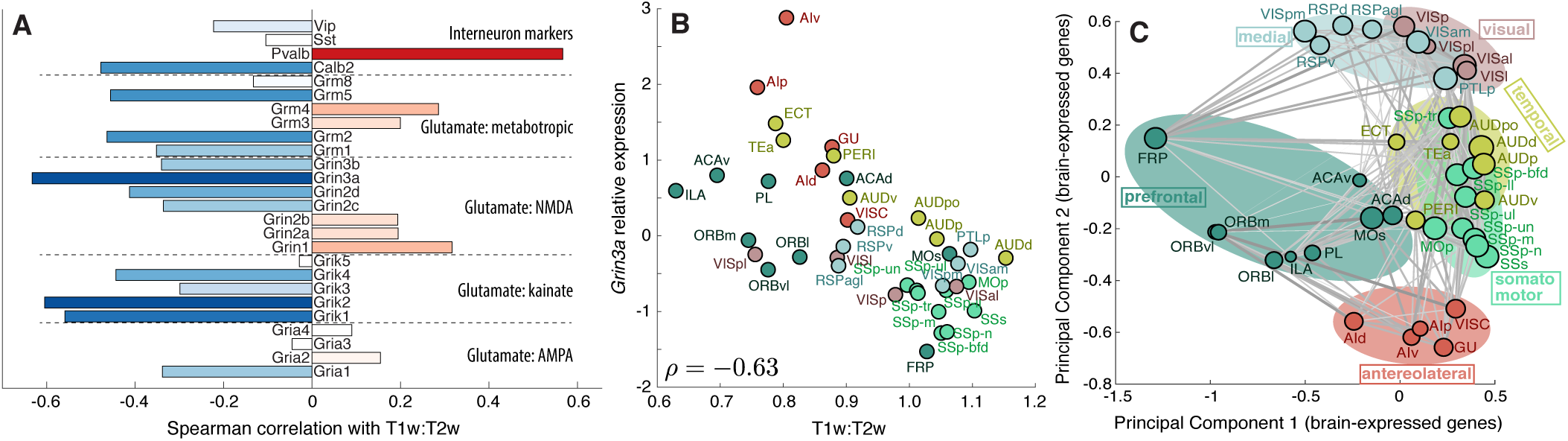
Transcriptional maps of glutamate receptor subunit and interneuron marker genes covary with the T1w:T2w map, and a principal components projection of cortical areas using their gene expression profiles organizes them into meaningful processing streams. **A** Spearman correlation coefficients, *ρ*, between T1w:T2w and the transcriptional maps of genes coding glutamate receptor subunits and interneuron cell-type markers. **B** Scatter plot of T1w:T2w versus *z*-score normalized levels of *Grin3a* transcription, *ρ* = −0.63 (*p*_corr_ = 5 × 10^−4^). **C** Projection of brain areas into the space of the two leading principal components of 1055 brain-expressed genes (52) places cortical areas with similar transcriptional profiles close in the space, yielding a functionally informative organization of cortical areas. Brain areas are shown as circles with radii scaled by T1w:T2w and (symmetrized) axonal projections (33) are annotated where possible.

To understand how T1w:T2w relates to dominant transcriptional gradients of a set of 1055 brain-expressed genes (52), we used principal components analysis (PCA) to estimate the most explanatory spatial maps of transcriptional variation (accounting for missing values using probabilistic PCA (55), see Supporting Information). The first PC of cortical transcription is significantly correlated with the T1w:T2w map, |*ρ*| = 0.53 (*p* = 6 × 10^−4^), as it is in human cortex, |*ρ*| = 0.81 (25) (the correlation with PC2 is weaker, |*ρ*| = 0.29, cf. Fig. S4). Figure 2C displays a projection of cortical areas into the space of the two leading PCs, placing areas with similar transcriptional profiles close to one another. This transcriptional organization of cortical areas clearly separates different functional processing streams and visually resembles parallel primary–transmodal hierarchies (22) that have recently been characterized in human rs-fMRI (18).

### Laminar specificity

Are large-scale cortical gradients driven by the specialization of specific cortical layers? We investigated this question using layer-specific maps of gene transcription (32) and interneuron density (29). T1w:T2w was estimated for each brain area by combining all cortical layers (values of T1w:T2w computed in layers 1-5 were highly correlated to this overall measurement, see Fig. S5). We first computed Spearman correlation coefficients, *ρ*, between T1w:T2w and cell densities of three types of interneurons (29) in each of five cortical layers: 1 (37 areas), 2/3 (37 areas), 4 (21 areas), 5 (36 areas), and 6 (35 areas). Results are plotted in Fig. 3A for each cortical layer (row) and cell type (column). Correcting across 15 (assumed independent) hypothesis tests (50)—each cell type in each cortical layer— we found a significant positive correlation between T1w:T2w and PV cell density in layer 5, *ρ* = 0.52 (*p*_corr_ = 7 × 10^−3^), and a negative correlation of SST cell density with T1w:T2w in layer 2/3, *ρ* = −0.62 (*p*_corr_ = 4 × 10^−4^) and layer 6, *ρ* = −0.65 (*p*_corr_ = 4 × 10^−4^). VIP cell density did not exhibit a significant correlation to T1w:T2w in any individual cortical layer.

**Fig. 3.**
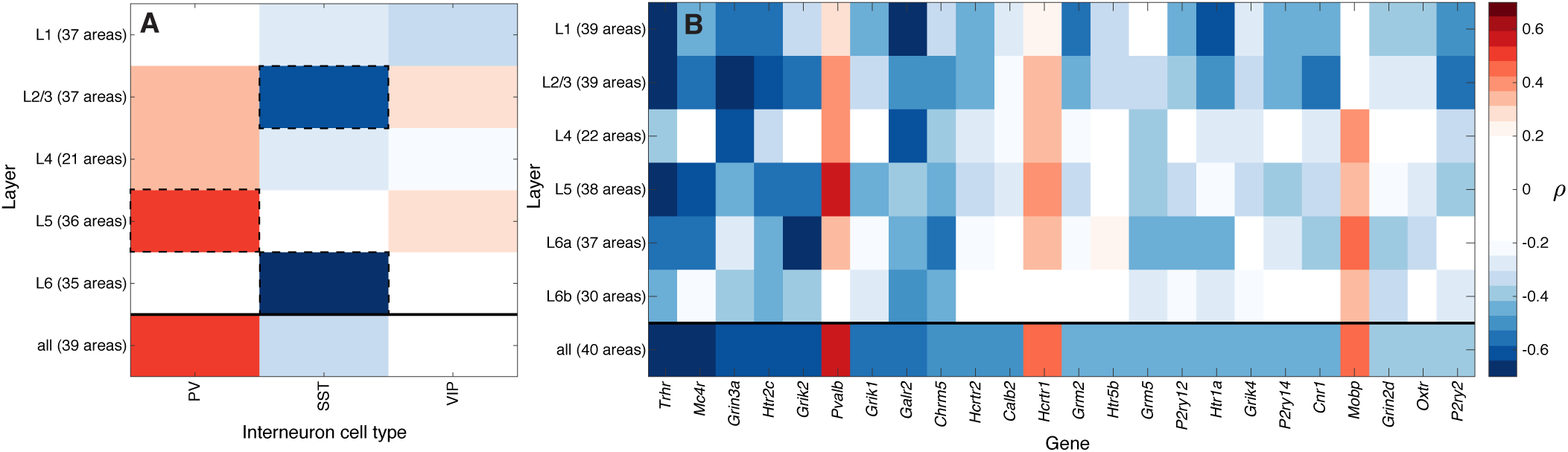
Cell densities and gene expression exhibit distinctive laminar patterns of covariation with T1w:T2w across the mouse cortex. Spearman correlation coefficients, *ρ*, of T1w:T2w are plotted as color for: **A** cell density of three types of interneurons (29), and **B** transcriptional level of a given gene (column) in a given cortical layer (row). Overall correlations, obtained from combining all layers of each area, are shown in the bottom row of each plot. Transcriptional levels (32) are shown for the 24 genes with significant overall correlations to T1w:T2w, ordered by their overall correlation to T1w:T2w (results for all 86 brain-related genes are in Fig. S7).

We next investigated gene-expression patterns in cortical layers 1, 2/3, 4, 5, 6a, and 6b (32) across the 24 genes identified above to be significantly correlated to T1w:T2w, shown in Fig. 3B (results for all 86 genes are in Fig. S7). Consistent with *Pvalb* expression as a marker of PV cell density, the two measurements exhibit a similar laminar pattern of T1w:T2w correlations (with the strongest correlation in layer 5, cf. Fig. 3A) and a high overall correlation (*ρ* = 0.82, see Supplementary Information). For a given gene, the direction of correlation between its expression and T1w:T2w is mostly consistent across cortical layers. Some genes display an association with T1w:T2w in all individual layers (e.g., *Trhr, Grin3a*, and *Htr2c*), while other genes show an overall correlation with T1w:T2w that is restricted to specific cortical layers (e.g., the positive correlation with *Mobp* is driven by layer 4 and infragranular layers). These cell density and gene expression results demonstrate that macroscopic gradients of areal specialization are not dominated by particular cortical layers; all layers exhibit macroscopic gradients of areal specialization, for distinct microcircuit properties.

### A common hierarchical gradient

The variation of T1w:T2w mirrors the large-scale variation of microstructural properties along a putative functional hierarchy of cortical areas. To more directly understand relationships between the diverse cortical gradients characterized above, we combined representative measurements from each data type: gene expression, intracortical axonal connectivity, T1w:T2w, and interneuron cell density. These properties were visualized together by plotting the cortical variation of each measurement type with a distinct color map, as shown in Fig. 4. Measurements (columns) with highly correlated variation across cortical areas (rows) were placed close to each other using linkage clustering (see Supporting Information). A common gradient emerges from the covariation of these diverse measurements of cortical structure, estimated as the first principal component of the data and used to reorder the rows of Fig. 4. This consensus gradient orders areas along a putative functional hierarchy, from primary somatosensory through to integrative prefrontal areas. Each individual measurement either increases or decreases along this consensus gradient, forming the two anticorrelated clusters shown in Fig. 4.

**Fig. 4.**
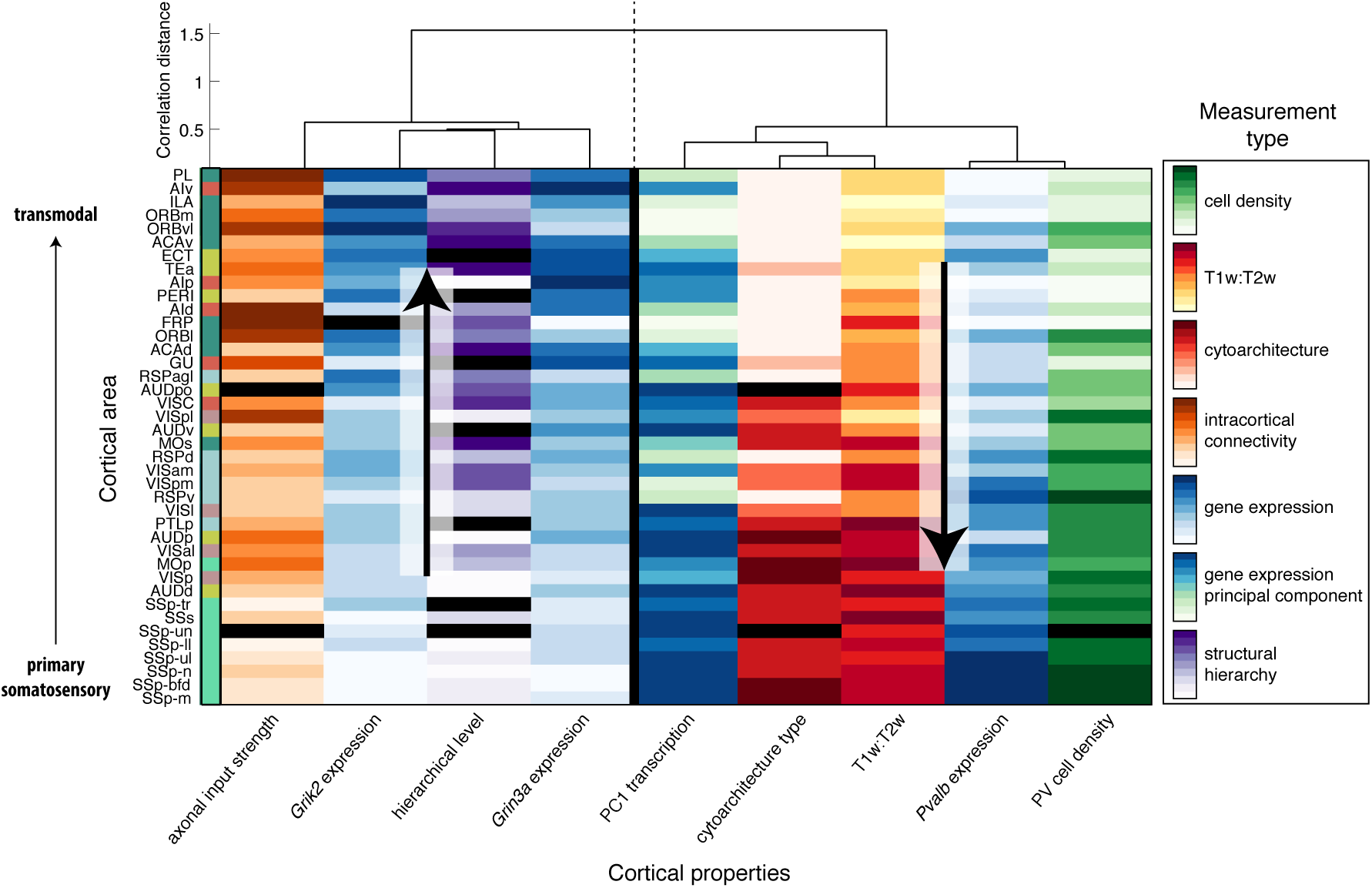
Diverse measurements of cortical areas share a common gradient of variation. Cortical areas form rows, and measurements form columns, with different color maps used to code different datasets(from low values, light, to high values, dark), as labeled: density of PV neurons (29) (green), T1w:T2w (53) (yellow/red), cytoarchitecture (37) (red), weighted in-degree, *k*^*w*^, of normalized axonal projection density (33) (orange), expression of three selected genes: *Grin3a, Grik2*, and *Pvalb* (32) (blue) and the first principal component of brain-related genes (green-blue), and estimated hierarchical level (30) (purple). Areas (rows) are ordered according to the first principal component of this multimodal matrix of cortical properties and are colored according to their membership to one of six connectivity-based groupings (30) (cf. Fig. 1B). Measurements (columns) are ordered according to average linkage clustering on correlation distances according to the annotated dendrogram. Measurements cluster into two groups that either increase (left cluster) or decrease (right cluster) along a candidate functional hierarchy, from primary somatosensory through to transmodal prefrontal areas. Missing data are shown as black rectangles.

To investigate whether ordering areas by T1w:T2w or the structural hierarchical ordering of Harris et al. (30) better explains the multimodal cortical gradients, we compared how each correlates with the properties shown in Fig. 4 (across the 35 cortical areas common to both measurements). Spearman correlation coefficients were similar in magnitude between T1w:T2w and hierarchical level, and all but one property (PV cell density) were more strongly correlated with T1w:T2w than with hierarchical level (see Fig. S8).

### Mouse–human consistency between T1w:T2w and gene transcriptional gradients

T1w:T2w follows dominant transcriptional gradients in mouse and human cortex (25), but are the gradients of specific brain-related genes conserved between the two species? To investigate this, we compared the correlation between T1w:T2w and the transcription map of a given gene in mouse cortex, *ρ*_*m*_, and for the human ortholog of that gene in human cortex, *ρ*_*h*_ (25), repeating the calculation the 70 of our 86 brain-related genes analyzed that have human orthologs and could be matched between the two datasets. Interspecies consistency in transcriptional gradients is reflected by a correspondence between the independent measurements of *ρ*_*m*_ and *ρ*_*h*_, which we measured as a correlation across genes as *ρ*_*mh*_. Although we have been careful to develop and apply rigorous quality control criteria, the AMBA (32) and Allen Human Brain Atlas (AHBA) (56) can be noisy at level of individual experiments; consequently, weak correlations with T1w:T2w in either human (low *ρ*_*h*_) or mouse (low *ρ*_*m*_) should not be interpreted as an absence of a relationship as much as strong correlations can be interpreted as evidence for a relationship.

As shown in Fig. 5, we find significant interspecies correspondence, *ρ*_*mh*_ = 0.44 (*p* = 1 × 10^−4^). The agreement is striking given measurement noise, vast differences in spatial scale, and distinct measurement modalities between mouse (high-throughput *in situ* hybridization) and human (post-mortem microarray from six adults). Two of the genes with the strongest correlations with T1w:T2w in mouse cortex exhibit a similar variation in human cortex: *Grin3a*/*GRIN3A (ρ*_*m*_ = – 0.63, *ρ*_*h*_ = – 0.65) and *Pvalb*/*PVALB* (0.57, 0.70). A range of other key genes exhibit strong interspecies consistency, including the interneuron marker *Calb2* (–0.48, *–*0.45), the oxytosin receptor gene, *Oxtr* (–0.41, *–*0.48), glutamate receptor genes *Grik1* (–0.56, *–*0.54), *Grik2* (–0.60, *–*0.52), and *Grik4* (–0.44, *–*0.33), and myelin marker genes *Mobp* (0.43, 0.41) and *Mbp* (0.34, 0.45). Significant mouse–human correspondence was not limited to the brain-related genes shown in Fig. 5, but was reproduced for: (i) all 2951 genes, *ρ*_*mh*_ = 0.25 (*p* = 8 ×10^−42^); (ii) 806 brain-expressed genes (52), *ρ*_*mh*_ = 0.31 (*p* = 2 × 10^−19^); (iii) 60 astrocyte-enriched genes, *ρ*_*mh*_ = 0.38 (*p* = 3 × 10^−3^); (iv) 143 neuron-enriched genes, *ρ*_*mh*_ = 0.40 (*p* = 6 × 10^−7^); and (v) 41 ogligodendrocyte-enriched genes, *ρ*_*mh*_ = 0.65 (*p* = 7 × 10^−6^) (57). Consistent with the enhancement of a meaningful signal, mouse–human correspondence increased as progressively more stringent quality-control criteria were applied to the mouse gene-expression data (see Fig. S9).

**Fig. 5.**
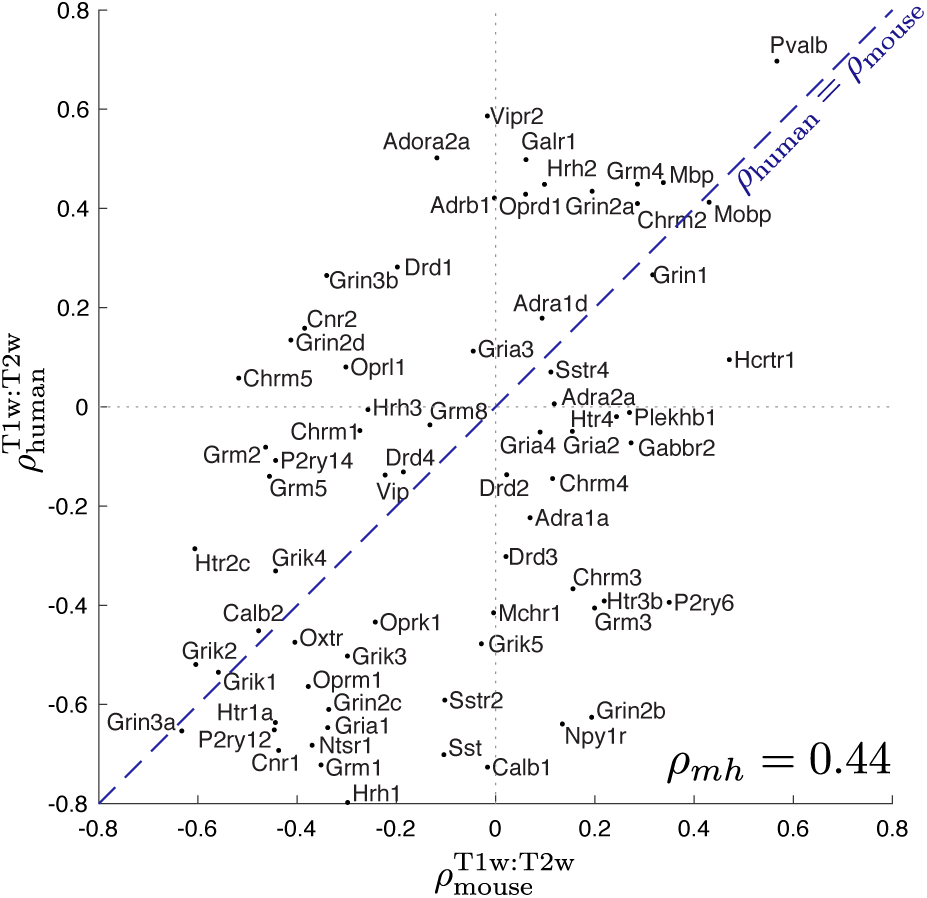
Mouse–human consistency of T1w:T2w and transcriptional gradients of brain-related genes. We plot the correlation between T1w:T2w and transcription levels in mouse, 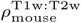 (horizontal axis) and human, 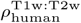 (vertical axis) for 70 brain-related genes. The equality line, *ρ*_*m*_ = *ρ*_*h*_, is shown dashed blue. Transcriptional maps of key brain-related genes vary similarly with T1w:T2w in mouse and human cortex, *ρ*_*mh*_ = 0.44 (*p* = 1 × 10^−4^).

## Discussion

Macroscale spatial variations in the makeup of cortical microcircuits may reflect a biological substrate of functional specialization that enables efficient processing and integration of diverse sensory information (5, 7, 10, 20, 21, 25) across multiple timescales (58–60). Here we show that these large-scale hierarchical gradients observed in macaque and human cortex (25) match those in mouse, which displays an under-appreciated level of interareal heterogeneity. Circumventing the need to define an interspecies homology (61), we used the noninvasive MRI contrast map, T1w:T2w, as a common spatial reference for interspecies comparison; T1w:T2w varies similarly in both mouse and human cortex with the dominant cortical map of transcriptional variation (estimated using PCA) and with specific transcriptional maps of genes encoding synaptic receptors and neuronal cell types. Using high-resolution invasive measurements available in mouse, we report new connections between T1w:T2w and cell densities, laminar-specific gene expression, and interareal axonal connectivity. Our results provide new understanding of the anatomical underpinning of hierarchical functional specialization, and demonstrate the usefulness of using noninvasive measurements like T1w:T2w to index large-scale gradients across species. While the complexity of cortical diversity clearly cannot be indexed by a single measurement like T1w:T2w, the covariation of diverse aspects of microcircuit architecture along a common spatial map, and its inter-species correspondence, supports the existence of stereotypical anatomical properties that support hierarchical specialization in mammalian brains.

The degree of interareal variation of microstructural properties in mouse (8, 9) is far less pronounced than in the highly differentiated primate cortex (7, 10, 62). For example, in rhesus monkey, the physiology and morphology of layer 3 pyramidal neurons (26) and glutamatergic synaptic structure (27) exhibit strong interareal differences between V1 and frontal cortex, properties that are strikingly homogeneous by comparison between homologous areas of mouse cortex. While the mouse cortex exhibits hierarchical feedforward-feedback projection patterns in specific (e.g., visual) processing streams (63), with a corresponding variation in microstructure (64), interareal laminar projections across the whole cortex appear to be more stereotyped in macaque (65) than mouse (30). Different functional modules of the mouse cortex fit a hierarchical organization to differing extents, with a low hierarchy score for the prefrontal module (0.03) and higher scores for visual (0.33) and temporal (0.51) modules, where a maximum score of 1 corresponds to an ideal hierarchy (30). Across the whole cortex, mouse brain areas fit relatively poorly into a global hierarchical organization, attaining a hierarchy score of just 0.126, although this index is yet to be calculated in macaque, preventing direct interspecies comparison. Consistent with a lesser degree of differentiation in mouse cortex, here we report consistently weaker correlations of T1w:T2w with other spatial maps relative to macaque and human, for hierarchical level (*ρ*_macaque_ = −0.76, *ρ*_mouse_ = −0.29), cytoarchitectural type (*τ*_macaque_ = 0.87, *τ*_mouse_ = 0.51), and the leading principal component of gene transcription (|*ρ*_mouse_| = 0.53, |*ρ*_human_| = 0.81) (25). A small number of genes exhibit mouse-human differences in the direction of their expression gradients with T1w:T2w (Fig. 5), but in most of these cases the correlation is weak in either mouse or human and may be attributable to measurement noise in the atlas-based expression data used here. However, our analysis flags candidate genes that further investigation may reveal to show robust mouse-human differences in the direction of expression gradients with T1w:T2w, including NMDA receptor signaling genes *Grin2b*/*GRIN2B (ρ*_*m*_ = 0.19, *ρ*_*h*_ = −0.63), *Grin2d*/*GRIN2D (ρ*_*m*_ = −0.41, *ρ*_*h*_ = 0.13), and *Grin3b*/*GRIN3B (ρ*_*m*_ = −0.34, *ρ*_*h*_ = 0.26).

As well as interspecies differences, our results also highlight an under-appreciated spatial dimension of cortical specialization in mouse that matches cortical gradients in primate. The consistency of transcriptional maps of 70 receptor subunit and cell-type marker genes with the common reference map, T1w:T2w, in mouse and human (*ρ*_*mh*_ = 0.44, *p* = 1 × 10^−4^) is striking given vastly different spatial scales and major differences in expression measurement between mouse (high-throughput *in situ* hybridization (32)) and human (microarray data from six post-mortem subjects (56)). In contrast to findings above, in which we found a generally weaker relationship of cortical gradients to T1w:T2w in mouse relative to primate, these transcriptional gradients were comparable in magnitude between the two species. A two-dimensional projection of mouse cortical areas based on their transcriptional signatures across brain-expressed genes (Fig. 2C), distinguishes somatomotor, auditory, and visual processing streams from antereolateral and prefrontal areas and yields an organization analogous to parallel processing streams in low-dimensional embeddings of human fMRI correlation networks (18, 66).

T1w:T2w is commonly interpreted as a marker of gray-matter myelin content (45), although both T1- and T2-weighted images are sensitive to a wide range of microstructural properties (46). Our results provide transcriptional evidence for a connection between T1w:T2w and myelin (67), demonstrating a significant relationship with the expression of *Mobp* and other oligodendrocyte-enriched genes (which also display high mouse–human correspondence, *ρ*_*mh*_ = 0.65). Interpreting T1w:T2w as a marker of relative intracortical myelination is consistent with the increase in T1w:T2w across areas with increasing laminar differentiation, which is associated with myelin content (4, 5, 48). Axonal myelination improves transmission speeds (68) and prevents the formation of new synaptic connections (69, 70), properties that are consistent with the fast, ‘hard-wired’ computations in heavily myelinated and eulaminar somatosensory areas relative to more plastic and adaptive agranular prefrontal areas (3, 5). While these diverse gradients may reflect associated differences in myelination, other gradients reported here cannot be linked straightforwardly to relative myelination levels, such as the strong transcriptional variation of *Grin3a* and *Calb2* with T1w:T2w. The convergence of multimodal cortical gradients may therefore reflect deeper organizational mechanisms acting in concert, perhaps through development (51). Our results are consistent with existing models of mammalian cortical organization based on systematic structural variation as a core organizing principle (48); in this context, the convergence of diverse gradients found here could be used to predict laminar patterns of interareal connectivity that do not rely on a global hierarchical representation of cortical areas. To understand the functional importance of the dominant cortical gradient reported here, further work is needed to explain how characteristic differences in synaptic structure and inhibitory control may allow for more flexible behavior at the level of information processing within local microcircuits (3, 48).

As more is learned about how variations in cellular and synaptic microstructure shape functional specialization in the cortex, it will be important to complement this understanding with new mathematical models of brain dynamics that are properly constrained by the breadth and spatial detail of new datasets (29, 59, 71, 72). Such approaches will allow theory and experiment to develop in tandem, with physiologically-constrained mathematical models making functional predictions that can be tested experimentally and used to refine the models. Understanding how individual cortical areas—each treated as a local computational unit (73) with distinctive dynamical properties—communicate coherently on a whole-brain scale, may aid motivation for the next generation of brain-inspired machine-learning algorithms (74). Our findings offer guidance for the development of dynamical and functional models of large-scale cortical circuits in mammalian species.

## ACKNOWLEDGEMENTS

We would like to thank Alex Fornito for his detailed and insightful comments on the manuscript. B.D.F is supported by the National Health and Medical Research Council 1089718. J.D.M. is supported by NIMH grant R01 MH112746. V.Z. is supported by the SNSF AMBIZIONE PZ00P3_173984. X.J.W. is supported by NIMH grant R01MH062349, Simons Collaborative Global Brain (SCGB) Program Grant 543057SPI, and ONR N00014-17-1-2041.

