## Supporting Information for "Multimodal gradients across mouse cortex"

### Data

**Cortical parcellation.** The 40 mouse cortical areas analyzed here are from the Allen Reference Atlas (ARA) (1), and labeled (where possible) according to the following grouping from Harris et al. (2):
***Somatomotor***: 'MOp' (Primary motor area), 'SSp-n' (Primary somatosensory area, nose), 'SSp-bfd' (Primary somatosensory area, barrel field), 'SSp-ll' (Primary somatosensory area, lower limb), 'SSp-m' (Primary somatosensory area, mouth), 'SSp-ul' (Primary somatosensory area, upper limb), 'SSp-tr' (Primary somatosensory area, trunk), 'SSp-un' (Primary somatosensory area, unassigned), 'SSs' (Supplemental somatosensory area).
***Medial***: 'PTLp' (Posterior parietal association areas), 'VISam' (Anteromedial visual area), 'VISpm' (Posteromedial visual area), 'RSPagl' (Retrosplenial area, lateral agranular part), 'RSPd' (Retrosplenial area, dorsal part), 'RSPv' (Retrosplenial area, ventral part).
***Temporal***: 'AUDd' (Dorsal auditory area), 'AUDp' (Primary auditory area), 'AUDpo' (Posterior auditory area), 'AUDv' (Ventral auditory area), 'TEa' (Temporal association areas), 'PERI' (Perirhinal area), 'ECT' (Ectorhinal area).
***Visual***: 'VISal' (Anterolateral visual area), 'VISl' (Lateral visual area), 'VISp' (Primary visual area), 'VISpl' (Posterolateral visual area).
***Anterolateral***: 'GU' (Gustatory areas), 'VISC' (Visceral area), 'AId' (Agranular insular area, dorsal part), 'AIp' (Agranular insular area, posterior part), 'AIv' (Agranular insular area, ventral part).
***Prefrontal***: 'FRP' (Frontal pole, cerebral cortex), 'MOs' (Secondary motor area), 'ACAv' (Anterior cingulate area, ventral part), 'ACAd' (Anterior cingulate area, dorsal part), 'PL' (Prelimbic area), 'ILA' (Infralimbic area), 'ORBl' (Orbital area, lateral part), 'ORBm' (Orbital area, medial part), 'ORBvl' (Orbital area, ventrolateral part).
The cortical areas analyzed here differ slightly from those analyzed by Harris et al. (2): (i) 'VISrl' and 'VISa' are grouped as 'PTLp' (and assigned to the 'medial' grouping), and (ii) 'VISli' (laterointermediate area) and 'VISpor' (postrhinal area) are excluded, as they are not in Oh et al. (3).

**Spatial coordinates.** Three-dimensional spatial coordinates of each brain region were computed as the mean of right-hemisphere cortical masks obtained for each individual region from the Allen SDK (2015)[1]. Reference space masks for each brain area were retrieved in the 25$\mu$m grid of the Allen Common Coordinate Framework (CCF v3), including only voxels in the right hemisphere by including only voxels with a $z$-coordinate $> 227$.

**Structural MRI.** T1w:T2w data was obtained from the scalable brain atlas (4) in Waxholm space (5). Images were rescaled to 25 $\mu$m isotropic voxel spacing and normalized to the Allen Reference Atlas (ARA) in CCFv3 using linear affine and non-linear greedy deformations with Advance Normalization Tools (ANTs)[2]. This yielded a high-quality normalization that is precise to a very high spatial resolution, as demonstrated in Figs 1, 2, and 3. Accordingly, variability in T1w:T2w measurements due to errors in transforming between Waxholm and CCFv3 space are considered to be negligible.
Although we focus on T1w:T2w throughout this work, T1w is strongly (negatively) correlated to T2w across the 40 mouse cortical areas analyzed, $\rho = -0.77$ ($p = 1 \times 10^{-7}$) and T1w:T2w is strongly correlated to both T1w ($\rho = 0.90$) and T2w ($\rho = -0.96$). Taking the ratio, as T1w:T2w, affects results minimally compared to using either T1w or T2w on their own. A high correspondence between T1w and T1w:T2w has also been noted in human (6, 7).

**Cell densities.** Cell counts were obtained for all 40 cortical areas from the single-cell annotation *CUBIC-Atlas* (8). Counts were converted to (volume) densities by dividing by the volume of each area, estimated as the number of labeled voxels in the 25$\mu$m-gridded Allen CCF v3. T1w:T2w is not significantly correlated to variation in cell density, $\rho = 0.21$ ($p = 0.2$).
Our main analysis of T1w:T2w and neuron density is taken from Erö et al. (9), based on the supplementary data table of cell volume densities (mm$^{-3}$). A total of 39 cortical areas matched by name to our Allen Reference Atlas cortical parcellation (all areas except for SSp-un). In each area, cell density data was available for each of: cells, neurons, glia, excitatory cells, inhibitory cells, modulatory cells, astrocytes, oligodendrocytes, and microglia. T1w:T2w is not significantly correlated to variations in neuron density, $\rho = -0.08$ ($p = 0.63$, across 39 matching cortical areas), or cell density, $\rho = 0.25$ ($p = 0.1$). Note

[1] http://alleninstitute.github.io/AllenSDK/
[2] stnava.github.io/ANTs/

the consistency between the correlation to cell density measured with this dataset ($\rho = 0.25$) and that of Murakami et al. (8) ($\rho = 0.21$).

To check for consistency with the data of Herculano-Houzel et al. (10), we took neuron area densities ($N$/mm$^2$) from Table 1 (10) across each of sixteen cortical areas of the Franklin and Paxinos mouse brain atlas (11). Values were then matched manually to ARA areas (1) by name, as listed in Table 1. A T1w:T2w value was then estimated for each Franklin-Paxinos area as the mean T1w:T2w across matched ARA areas. As we found using more comprehensive data above (9), T1w:T2w is not significantly correlated to neuron density, $\rho = 0.20$ ($p = 0.5$).

Interneuron cell densities were measured by *qBrain*, quantitative whole-brain mapping of distributions of fluorescently labeled neural cell types (12). The cell counting and distribution mapping platform involves automated imaging by serial two-photon tomography, followed by a machine-learning based analysis pipeline; results are accessible via the web portal: http://mouse.brainarchitecture.org/cellcounts/ost/. We used data provided directly from the authors, which provide agglomerated information across repeats of experiments in the ARA. For each brain region, we took the mean cell density (across 10 repeat experiments) for each of parvalbumin-containing (PV), somatostatin-containing (SST), and vasoactive intestinal peptide-containing (VIP) cells. We analyzed data from bulk cortical areas as well as the same data delineated by cortical layer.

**Gene expression.** Gene expression data was obtained from the Allen Mouse Brain Atlas (AMBA) (13) using the Allen Software Development Kit (SDK)[3]. Gene expression data in the AMBA is measured using *in situ* hybridization from: (i) sagittal section experiments with high genome coverage, and (ii) coronal section replications for approximately 3 500 genes with restricted expression patterns in the brain (13). Transcriptional levels across a macroscopic cortical area was summarized as the 'expression energy' (the mean ISH intensity across voxels of that brain area) (13, 14).

Previous studies have represented the transcriptional profile of a brain area across a large number of genes (15, 16). In such a representation, even if an individual gene contributes a highly noisy datapoint, meaningful patterns can emerge from the aggregate contribution of large sets of genes (and captured using, for example, enrichment analysis at the level of functional categories (17)). By contrast, in this work we aimed to interpret the spatial expression maps of individual genes, which required us to develop and apply more stringent quality-control criteria than used in previous work. Genes included in our analysis analysis had either: (i) coronal section data available, or (ii) multiple section datasets exhibiting high agreement ($r \geq 0.5$ across mouse cortical areas). These criteria represent a compromise between maintaining data quality and maximizing gene coverage: of the 19 419 genes in the AMBA, 4181 fulfil these criteria. These 4181 genes are enriched in brain-related function, by virtue of both the choice of genes measured using coronal section data in AMBC, and through our criterion of requiring consistent cortical expression patterns across multiple experiments. Where multiple section datasets were available for the same gene, they were combined by first $z$-scoring expression data taken across areas, and then computing the expression value of a cortical area as the mean across $z$-scored section datasets. Below we demonstrate that mouse–human correspondence of transcriptional gradients (with T1w:T2w) increases with the stringency of gene inclusion, supporting the proposition that the quality control procedures applied are concentrating meaningful signal in the data.

Layer-specific gene expression was also downloaded using the Allen SDK[4], by retrieving expression data for all child structures of the isocortex (structure 315 in the Allen Mouse Reference Atlas), and filtering by those containing 'layer' in their name, yielding a set of 297 cortical structures with layer-specificity, covering all 40 cortical areas listed above. In total, we retrieved gene expression data for 'layer 1' (39 areas), 'layer 2/3' (39 areas), 'layer 4' (22 areas), 'layer 5' (38 areas), 'layer 6a' (37 areas), and 'layer 6b' (30 areas).

**Custom gene sets.** To aid interpretation of our results, we analyzed a range of specific gene sets, which are described here. A list of brain-expressed genes was obtained from the gene expression database, GXD (18), by filtering on detected expression in 'brain' in wild-type mice of any age or assay type. This list contained 2421 genes, of which 2331 could be matched to the AMBA by gene symbol, and 1055 of these met our quality criteria. Genes with greater than a 10-fold enrichment in neurons, astrocytes, and ogligodendrocytes were labeled according to data from Cahoy et al. (19). After filtering genes to those meeting our quality criteria, we had three gene sets: (i) 171 (/251) neuron-enriched genes, (ii) 47 (/100) ogligodendrocyte-enriched genes, and (iii) 71 (/157) astrocyte-enriched genes.

***Brain-related genes.*** The following list of brain-related genes is based on a list of receptor subunit genes from Janusonis et al. (20). We also added the four 'most abundant mRNAs in myelin': *Mbp*, *Fth1*, *Plekhb1*, and *Mobp* (21), as well as some interneuron cell-type markers, as listed below. Genes that are missing from the AMBA are shown red and genes that did not meet our quality control criteria are shown blue. 120 genes on this list are contained in the AMBA, and 86 of these met our quality criteria.

[3]©2010 Allen Institute for Brain Science. Allen SDK. Available from: http://alleninstitute.github.io/AllenSDK/
[4]©2010 Allen Institute for Brain Science. Allen SDK. Available from: http://alleninstitute.github.io/AllenSDK/

***Receptors***
**Adrenergic receptor (5/9)**: *Adra1a*, *Adra1b*, *Adra1d*, *Adra2a*, *Adra2b*, *Adra2c*, *Adrb1*,*Adrb2*, *Adrb3*.
**Adenosine receptor (1/4)**: *Adora1*, *Adora2a*, *Adora2b*, *Adora3*.
**Cannabinoid receptor (2/2)**: *Cnr1*, *Cnr2*.
**Cholinergic receptor (5/5)**: *Chrm1*, *Chrm2*, *Chrm3*, *Chrm4*, *Chrm5*.
**Dopamine receptor (4/5)**: *Drd1*, *Drd2*, *Drd3*, *Drd4*, *Drd5*.
**GABA receptor (1/2)**: *Gabbr1*, *Gabbr2*.
**Galanin receptor (2/3)**: *Galr1*, *Galr2*, *Galr3*.
**Glutamate receptor–AMPA (4/4)**: *Gria1*, *Gria2*, *Gria3*, *Gria4*.
**Glutamate receptor–kainate (5/5)**: *Grik1*, *Grik2*, *Grik3*, *Grik4*, *Grik5*.
**Glutamate receptor–NMDA (7/7)**: *Grin1*, *Grin2a*, *Grin2b*, *Grin2c*, *Grin2d*, *Grin3a*, *Grin3b*.
**Glutamate receptor–metabotropic (6/8)**: *Grm1*, *Grm2*, *Grm3*, *Grm4*, *Grm5*, *Grm6*, *Grm7*, *Grm8*.
**Histamine receptor (3/4)**: *Hrh1*, *Hrh2*, *Hrh3*, *Hrh4*.
**Hypocretin receptor (2)**: *Hcrtr1*, *Hcrtr2*.
**Melanocortin receptor (2/3)**: *Mc1r*, *Mc3r*, *Mc4r*.
**Melatonin-concentrating hormone (1)**: *Mchr1*.
**Neuropeptide Y receptor (1/5)**: *Npy1r*, *Npy2r*, *Npy4r*, *Npy5r*, *Npy6r*.
**Neurotensin receptor (1/2)**: *Ntsr1*, *Ntsr2*.
**Nociceptin receptor (1/1)**: *Oprl1*.
**Opioid receptor (3/3)**: *Oprm1*, *Oprd1*, *Oprk1*.
**Oxytosin receptor (1/1)**: *Oxtr*.
**Purine receptor (5/10)**: *P2ry1*, *P2ry2*, *P2ry4*, *P2ry6*, *P2ry10*, *P2ry12*, *P2ry13*, *P2ry14*, *P2rx1*.
**Serotonin receptor (8/14)**: *Htr1a*, *Htr1b*, *Htr1d*, *Htr1e*, *Htr1f*, *Htr2a*, *Htr2b*, *Htr2c*, *Htr3a*, *Htr3b*, *Htr4*, *Htr5a*, *Htr5b*, *Htr6*, *Htr7*.
**Somatostatin receptor (2/5)**: *Sstr1*, *Sstr2*, *Sstr3*, *Sstr4*, *Sstr5*.
**Tachykinin receptor (2/3)**: *Tacr1*, *Tacr2*, *Tacr3*.
**Thyrotropin releasing hormone receptor (1/2)**: *Trhr*, *Trhr2*.
**Vasopressin receptor (2/3)**: *Avpr1a*, *Avpr1b*, *Avpr2*.
**VIP receptor (1/2)**: *Vipr1*, *Vipr2*.

***Cell-type markers***
**Parvalbumin (1/1)**: *Pvalb*.
**Somatostatin (1/1)**: *Sst*.
**Calbindin (2/2)**: *Calb1*, *Calb2*.
**Vasoactive intestinal polypeptide (1/1)**: *Vip*.
**Myelin markers (3/4)**: *Mbp*, *Mobp*, *Plekhb1*, *Fth1*.

**Intracortical axonal connectivity.** Intracortical axonal connectivity data are based on 469 anterograde viral microinjection experiments in C57BL/6J male mice at age P56, obtained from the Allen Mouse Brain Connectivity Atlas (AMBCA) (3). Connection strengths and $p$-values were computed from a linear regression model across the whole mouse brain (3). We analyzed right-hemisphere ipsilateral intracortical axonal connectivity across 38 of our 40 cortical areas that matched those used by Oh et al. (3). Normalized connection density (NCD) was used a measure of edge weight, measuring the fraction of infected volume in the target region resulting from infection of a unit voxel of the source region, as used previously (16).
Alternate intracortical connectivity data are available from Ypma and Bullmore (22) and Gamanut et al. (23). Revised edge weight estimates of Ypma and Bullmore (22) (from a re-analysis of AMBCA data), did not yield any statistically significant correlations between T1w:T2w and $k^w_{\mathrm{in}}$ or $k^w_{\mathrm{out}}$. Intracortical axonal connectivity data collected independently by Gamanut et al. (23) are not registered to the ARA, so could not easily be matched to the cortical parcellation used here and were therefore not analyzed.

**Cytoarchitecture categorization.** A cytoarchitectonic classification of 38 cortical areas was obtained from Figure 1 of Goulas et al. (24). Labels denote distinct cortical types on an ordinal scale, ranging from 1 (less eulaminated) through to 4 (more eulaminated). Intermediate values denote areas that exhibit substantial within-area heterogeneity, and occur for four areas: VISam, VISpm, VISl, and VISpl (labeled as 2.5—a combination of types 2 and 3).

**Human gene expression.** To compare how gene-expression gradients relate to T1w:T2w between mouse and human cortex (Fig. 5), we used correlation values from Burt et al. (25); detailed methods are described therein. In brief, human gene expression maps were derived from the Allen Human Brain Atlas (AHBA) (26), from microarray measurements from the left hemisphere of six post-mortem subjects. The group-averaged ($N = 339$) T1w:T2w map was derived from the Human

Connectome Project (HCP) (27). Gene expression and T1w:T2w maps for the left cortical hemisphere were parcellated into 180 areas using the HCP's Multi-Modal Parcellation (MMP1.0) (28). Correlation values (Spearman $\rho$) between gene expression levels and T1w:T2w were computed over data from these 180 parcellated areas.
Human genes were mapped to mouse orthologs using data from Mouse Genome Informatics[5]. In this way, 12 265/16 039 human genes from Burt et al. (25) were successfully mapped to a unique mouse homolog Entrez ID (from a human gene symbol). These could be matched to Entrez IDs for 2951/4181 of genes in the AMBA that met our quality criteria.

### Methods

Code for reproducing our analyses is provided at `https://github.com/benfulcher/mouseInterneurons`. All analysis reported here was performed using a combination of python 2.7.13 and Matlab 2017b[6]. As distributions of cortical properties were frequently non-normally distributed, we computed Spearman rank correlations, $\rho$ to quantify relationships between two cortical properties. The only exception to this was for the categorical cytoarchitectural type, for which we used , or Kendall rank correlations, $\tau$, which better account for tied ranks in data (25). Correction for multiple hypothesis testing was achieved by controlling the false discovery rate at 5%, using the method of Benjamini and Hochberg (29). Correction was implemented in Matlab using the `mafdr` function and setting the `BHFDR` flag to `true`.

**Principal components analysis (PCA).** To understand the primary dimensions of brain-related transcriptional variance, we first constructed the 40 (region) $\times$ 1055 (brain-expressed gene) matrix, $X$. To reduce the effect of outliers in the data, we applied a sigmoidal transformation, $f(x) = [1+\exp(-\tilde{x})]^{-1}$ (where $\tilde{x}$ is a $z$-scored transformation of $x$) to each column, $x$, the result of which was then $z$-scored, as $\tilde{f}$, to ensure a zero mean and unit standard deviation for PCA. Because this matrix contained 79 missing values (1.8% of all values), we used probabilistic PCA (PPCA) (30) with an unrestricted Gaussian posterior for dimensionality reduction. PPCA yielded a set of components—spatial maps across 40 regions—ordered by their variance explained, that represent the projection of high-variance directions of combined gene expression into the space of brain regions. Specifically, for $n_{\text{comp}}$ components, we computed the weight matrix, $W$ ($n_{\text{gene}} \times n_{\text{comp}}$), that linearly projects the cortical transcription data, $X$ ($n_{\text{area}} \times n_{\text{gene}}$), into a space, $T$ ($n_{\text{area}} \times n_{\text{comp}}$), in which components of $T$ correspond to orthogonal spatial maps that progressively capture maximal variance of the full expression data, $X$, as $T = XW$. Note that our definition—the spatial projection of gene-expression data along principal components (columns of $T$)—differs from an alternative definition of a principal component as a set of loadings or weights across areas (columns of $W$). PPCA was implemented using the Matlab toolbox, *PCAMV*[7]. Similar results were obtained using an alternative method for performing PCA with missing values, alternating least squares algorithm for PCA (30), implemented natively in Matlab.

**Normalizing and clustering multimodal cortical gradients.** Visualizing multimodal properties of cortical areas was achieved by constructing a $n_{\text{areas}} \times n_{\text{properties}}$ matrix, $X$, shown in Fig. 4. Each column was normalized to the unit interval using an outlier-robust sigmoidal transformation (31). Pearson correlation distances were used to reorder columns according to average linkage clustering, implemented in Matlab using the `linkage` function. Rows were ordered according to a common multimodal gradient, computed as the first principal component of the normalized multimodal matrix, $X$, using PPCA (30) (as above).

### Supplementary Results

**Spatial embedding of T1w:T2w.** The T1w:T2w map can be well reconstructed as a linear gradient through three-dimensional space; using the three spatial coordinates of brain area centroids to predict T1w:T2w in a multilinear regression yields a high correlation, $\rho = 0.84$. This correlation is driven by an increase in T1w:T2w along the inferior–superior axis, $\rho = 0.53$ ($p = 5 \times 10^{-4}$). Correlations across the other individual spatial directions are weaker (anterior–posterior axis, $\rho = 0.17$ and the left–right axis, $\rho = -0.21$). Although T1w:T2w is spatially embedded, it demonstrates anatomical specificity beyond a monotonic spatial gradient, as shown in Fig. 1A.

**Validation of marker gene expression against cell-density measurements.** Access to independent measurements of both gene expression (13) and interneuron cell-type densities (12) allowed us to validate transcriptional markers against direct measurements of the cells that they index (32). We found a strong association between *Pvalb* and PV cell density across cortical areas, $\rho = 0.82$ ($p = 2 \times 10^{-8}$) [noting a strikingly strong correlation, $\rho = 0.95$ ($p \approx 0$), when using just the higher-quality coronal section data], a strong association also between VIP cell density and *Vip* gene expression, $\rho = 0.76$ ($p = 3 \times 10^{-7}$). The quality of the transcriptional data is supported by the strong correlation between independent measurements of interneuron

[5]World Wide Web (URLs: `www.informatics.jax.org/homology.shtml` and `ftp://ftp.informatics.jax.org/pub/reports/HOM_MouseHumanSequence.rpt`). [Data retrieved November 2014]
[6]Matlab is a product of The MathWorks, Natick, MA.
[7]downloaded from `http://users.ics.aalto.fi/alexilin/software/`

cell-type density, and these results suggest that transcriptional data can provide a highly accurate proxy for cell-type density. However, the relationship between SST cell density and *Sst* expression, $\rho = 0.24$ ($p = 0.1$), is not statistically significant. Even though all SST brain cells are GABAergic interneurons, levels of *Sst* expression decreases dramatically in adults despite SST+ cells remaining, with *Sst* expression regulated by activity (33); these factors may contribute to the lack of agreement between *Sst* expression and SST cell density. This result suggests that inferences of cell density from gene expression assays must in general be interpreted carefully due to the many (including time-varying) factors regulating gene transcription, on top of the many factors complicating straightforward inference of protein levels from mRNA (34).

**T1w:T2w as a myelin marker.** T1w:T2w is a candidate marker for gray-matter myelin (6, 35). Three of the four most abundant mRNAs in myelin—*Mbp*, *Plekhb1*, and *Mobp*—have transcriptional data that meet our quality criteria (*Fth1* was excluded) (21). T1w:T2w is correlated with the expression of these myelin-marker genes in the cortex, displaying a significant positive correlation with transcriptional levels of *Mobp*, $\rho = 0.43$ ($p_{\mathrm{corr}} = 0.018$), and non-significant positive correlations with *Mbp*, $\rho = 0.34$ ($p_{\mathrm{corr}} = 0.05$), and *Plekhb1*, $\rho = 0.27$ ($p_{\mathrm{corr}} = 0.09$). We also investigated whether genes enriched in ogligodendrocytes exhibited an increased correlation in expression profiles to T1w:T2w relative to neuron-enriched or astrocyte-enriched genes (taking a 10-fold enrichment threshold (19), yielding a set of 47 genes). The mean correlation between ogligodendrocyte enriched genes and T1w:T2w was modest, $\rho = 0.07 \pm 0.27$, but was significantly increased relative to 171 neuron-enriched genes (Wilcoxon signed-rank test, $p = 0.01$) and 71 astrocyte-enriched genes ($p = 0.007$). We next tested a set of 999 genes most abundant in myelin in cortex at age 6 months (21), of which 439 genes met our quality-control criteria. The mean correlation coefficient between T1w:T2w and these myelin-enriched genes is significantly increased relative to randomly selected genes (of those that met our quality-control criteria), $p < 1 \times 10^{-5}$ (permutation test with 100 000 repeats). Note that our 4181 random genes are not truly random, but are enriched in brain-related function due to our quality criteria for including genes. Together, these results provide transcriptional evidence for a relationship between T1w:T2w and myelin content (6, 35).

**Interspecies comparison of T1w:T2w–transcription relationships.** We investigated how interspecies consistency of T1w:T2w–transcriptional gradients, $\tilde{\rho}_{\mathrm{mh}}$, depends on data quality, hypothesizing that mouse-human consistency should increase when using higher-quality mouse expression data. We compared five criteria, which we expected to yield progressively accurate data and increase mouse-human correlations with T1w:T2w: (i) *'sagittal'*: using only sagittal section data (19 398 genes), (ii) *'all'* using data from both sagittal and coronal sections (19 419 genes), (iii) *'coronal'*: using only coronal section data (4064 genes), (iv) *'combination'*: the dual criteria described above that is used in this work (4181 genes), (v) *'replicated'*: using only genes that have been measured at least twice from pairs of section experiments that agree in their cortical expression profiles with a Pearson correlation of $r > 0.5$ (1311 genes). Results are shown in Fig. 9. As expected, mouse–human correspondance increases as increasingly stringent quality control criteria are applied: mouse–human correspondence is lowest when only sagittal data are used, is highest when only genes with multiple replicated section experiments are used. Note that this trend is driven both by increased data quality, and also from a progressive enrichment of brain-related genes following application of more stringent inclusion criteria.

**Outlook on continuum representations of the brain.** This work relies on sophisticated brain mapping experiments that measure the brain in unprecedented detail in standardized reference spaces (36), allowing new insights to be gained through combining diverse neuroscience data. In this way, cortical areas can be represented as a multimodal signature of diverse properties (Fig. 2) to understand cortical organizational principles that play out through the interplay of multiple processes. Future work using high-resolution spatial maps may move beyond the parcellation-based approach used here, towards a continuous data-driven representation of the brain in terms of continuous spatial gradients (37, 38). A continuum representation of the brain would also aid interspecies comparisons using the approach of taking a common measurement, such as T1w:T2w, as a common spatial reference map. Such an approach does not rely on the definition of a cross-species homology of areas and goes beyond simpler comparisons along e.g., rostro-caudal spatial axes (39).


1. H.-W. Dong. *The Allen reference atlas: A digital color brain atlas of the C57Bl/6J male mouse.* John Wiley & Sons Inc., Hoboken, NJ, US (2008).
2. J. A. Harris, S. Mihalas, K. E. Hirokawa, et al. The organization of intracortical connections by layer and cell class in the mouse brain. *bioRxiv* p. 292961 (2018).
3. S. W. Oh, J. A. Harris, L. Ng, et al. A mesoscale connectome of the mouse brain. *Nature* **508**, 207 (2014).
4. R. Bakker, P. Tiesinga, and R. Kötter. The Scalable Brain Atlas: Instant Web-Based Access to Public Brain Atlases and Related Content. *NeuroInf.* **13**, 353 (2015).
5. G. A. Johnson, A. Badea, J. Brandenburg, et al. Waxholm Space: An image-based reference for coordinating mouse brain research. *NeuroImage* **53**, 365 (2010).
6. J. Ritchie, S. P. Pantazatos, and L. French. Transcriptomic characterization of MRI contrast, focused on the T1-w/T2-w ratio in the cerebral cortex. *NeuroImage* **174**, 504 (2018).
7. J. M. Huntenburg, P.-L. Bazin, A. Goulas, et al. A Systematic Relationship Between Functional Connectivity and Intracortical Myelin in the Human Cerebral Cortex. *Cereb. Cortex* **27**, 981 (2017).
8. T. C. Murakami, T. Mano, S. Saikawa, et al. A three-dimensional single-cell-resolution whole-brain atlas using CUBIC-X expansion microscopy and tissue clearing. *Nat. Neurosci.* **21**, 625 (2018).
9. C. Erö, M.-O. Gewaltig, D. Keller, and H. Markram. A Cell Atlas for the Mouse Brain. *Front. Neuroinf.* **12**, e17727 (2018).
10. S. Herculano-Houzel, C. R. Watson, and G. Paxinos. Distribution of neurons in functional areas of the mouse cerebral cortex reveals quantitatively different cortical zones. *Front. Neuroanat.* **7**, 35 (2013).
11. K. B. L. Franklin and G. Paxinos. *The Mouse Brain in Stereotaxic Coordinates.* Elsevier Academic Press, San Diego, CA, 3rd edition (2007).
12. Y. Kim, G. R. Yang, K. Pradhan, et al. Brain-wide Maps Reveal Stereotyped Cell-Type-Based Cortical Architecture and Subcortical Sexual Dimorphism. *Cell* **171**, 456 (2017).
13. E. Lein, M. J. Hawrylycz, N. Ao, et al. Genome-wide atlas of gene expression in the adult mouse brain. *Nature* **445**, 168 (2006).
14. L. Ng, A. Bernard, C. Lau, et al. An anatomic gene expression atlas of the adult mouse brain. *Nat. Neurosci.* **12**, 356 (2009).
15. B. D. Fulcher and A. Fornito. A transcriptional signature of hub connectivity in the mouse connectome. *Proc. Natl. Acad. Sci. USA* **113**, 1435 (2016).
16. M. Rubinov, R. J. F. Ypma, C. Watson, and E. T. Bullmore. Wiring cost and topological participation of the mouse brain connectome. *Proc. Natl. Acad. Sci. USA* **112**, 10032 (2015).
17. M. Ashburner, C. A. Ball, J. A. Blake, et al. Gene Ontology: tool for the unification of biology. *Nat. Genet.* **25**, 25 (2000).

18. J. H. Finger, C. M. Smith, T. F. Hayamizu, et al. The mouse Gene Expression Database (GXD): 2017 update. *Nucl. Acid. Res.* **45**, D730 (2017).
19. J. D. Cahoy, B. Emery, A. Kaushal, et al. A transcriptome database for astrocytes, neurons, and oligodendrocytes: a new resource for understanding brain development and function. *J. Neurosci.* **28**, 264 (2008).
20. S. Janušonis. A receptor-based analysis of local ecosystems in the human brain. *BMC Neurosci.* **18**, 551 (2017).
21. S. Thakurela, A. Garding, R. B. Jung, et al. The transcriptome of mouse central nervous system myelin. *Sci. Rep.* **6**, 25828 (2016).
22. R. J. F. Ypma and E. T. Bullmore. Statistical Analysis of Tract-Tracing Experiments Demonstrates a Dense, Complex Cortical Network in the Mouse. *PLoS Comp. Biol.* **12**, e1005104 (2016).
23. R. Gămănuţ, H. Kennedy, Z. Toroczkai, et al. The Mouse Cortical Connectome, Characterized by an Ultra-Dense Cortical Graph, Maintains Specificity by Distinct Connectivity Profiles. *Neuron* **97**, 698 (2018).
24. A. Goulas, H. B. M. Uylings, and C. C. Hilgetag. Principles of ipsilateral and contralateral cortico-cortical connectivity in the mouse. *Brain. Struct. Funct.* **252**, 1 (2016).
25. J. B. Burt, M. Demirtas, W. J. Eckner, et al. Hierarchy of transcriptomic specialization across human cortex captured by structural neuroimaging topography. *Nat. Neurosci.* **27**, 889 (2018).
26. M. J. Hawrylycz, E. Lein, A. L. Guillozet-Bongaarts, et al. An anatomically comprehensive atlas of the adult human brain transcriptome. *Nature* **489**, 391 (2012).
27. D. C. Van Essen, S. M. Smith, D. M. Barch, et al. The WU-Minn Human Connectome Project: An overview. *NeuroImage* **80**, 62 (2013).
28. M. F. Glasser, T. S. Coalson, E. C. Robinson, et al. A multi-modal parcellation of human cerebral cortex. *Nature* **536**, 171 (2016).
29. Y. Benjamini and Y. Hochberg. Controlling the False Discovery Rate: A Practical and Powerful Approach to Multiple Testing. *J. Roy. Stat. Soc. B* **57**, 289 (1995).
30. A. Ilin and T. Raiko. Practical Approaches to Principal Component Analysis in the Presence of Missing Values. *J. Mach. Learn. Res.* **11**, 1957 (2010).
31. B. D. Fulcher, M. A. Little, and N. S. Jones. Highly comparative time-series analysis: the empirical structure of time series and their methods. *J. Roy. Soc. Interface* **10**, 20130048 (2013).
32. B. O. Mancarci, L. Toker, S. Tripathy, et al. Cross-Laboratory Analysis of Brain Cell Type Transcriptomes with Applications to Interpretation of Bulk Tissue Data . *eNeuro* **4**, e0212 (2017).
33. J. Urban-Ciecko and A. L. Barth. Somatostatin-expressing neurons in cortical networks. *Nat. Rev. Neurosci.* **17**, 401 (2016).
34. B. Schwanhäusser, D. Busse, N. Li, et al. Global quantification of mammalian gene expression control. *Nature Publishing Group* **473**, 337 (2011).
35. M. F. Glasser and D. C. Van Essen. Mapping Human Cortical Areas In Vivo Based on Myelin Content as Revealed by T1- and T2-Weighted MRI. *J. Neurosci.* **31**, 11597 (2011).
36. D. Fürth, T. Vaissière, O. Tzortzi, et al. An interactive framework for whole-brain maps at cellular resolution. *Nat. Neurosci.* **21**, 139 (2017).
37. J. M. Huntenburg, P.-L. Bazin, P. L. Bazin, and D. S. Margulies. Large-Scale Gradients in Human Cortical Organization. *TICS* **22**, 21 (2017).
38. K. V. Haak, A. F. Marquand, and C. Beckmann. Connectopic mapping with resting-state fMRI. *NeuroImage* (2017).
39. C. J. Charvet and B. L. Finlay. Evo-Devo and the Primate Isocortex: The Central Organizing Role of Intrinsic Gradients of Neurogenesis. *Brain Behav. Evol.* **84**, 81 (2014).

**Table 1.** Mapping of cortical areas of the Franklin-Paxinos mouse atlas (11) to the Allen Reference Atlas parcellation used here.

| Franklin Paxinos Area | Allen Reference Atlas Area(s) |
|---|---|
| Infralimbic | ILA |
| Cingulate | ACAd, ACAv |
| Retrosplenial | RSPd, RSPv, RSPagl |
| Parietal | PTLp |
| Motor | MOp, MOs |
| Frontal | FRP |
| V2M | VISam, VISpm |
| V1 | VISp |
| V2L | VISl |
| S1-limb | SSp-ll, SSp-ul |
| S1-face | SSp-m, SSp-n, SSp-bfd |
| S2 | SSs |
| Auditory | AUDp, AUDd, AUDv, AUDpo |
| Insula | AId, AIp, AIv |
| Orbital | ORBl, ORBm, ORBvl |
| Ectorhinal | ECT |

| Gene Name | Symbol | $\rho_{\mathrm{T1w:T2w}}$ | $p$ | $p_{\mathrm{corr}}$ |
|---|---|---|---|---|
| thyrotropin releasing hormone receptor | *Trhr* | -0.69 | 1.6e-06 | 0.00014 |
| melanocortin 4 receptor | *Mc4r* | -0.67 | 4.4e-06 | 0.00019 |
| glutamate receptor ionotropic, NMDA3A | *Grin3a* | -0.63 | 1.9e-05 | 0.00053 |
| 5-hydroxytryptamine (serotonin) receptor 2C | *Htr2c* | -0.61 | 4.8e-05 | 0.00083 |
| glutamate receptor, ionotropic, kainate 2 (beta 2) | *Grik2* | -0.60 | 4.6e-05 | 0.00083 |
| parvalbumin | *Pvalb* | 0.57 | 0.00018 | 0.0025 |
| glutamate receptor, ionotropic, kainate 1 | *Grik1* | -0.56 | 0.00023 | 0.0028 |
| galanin receptor 2 | *Galr2* | -0.54 | 0.00038 | 0.0041 |
| cholinergic receptor, muscarinic 5 | *Chrm5* | -0.52 | 0.00074 | 0.007 |
| hypocretin (orexin) receptor 2 | *Hcrtr2* | -0.49 | 0.0015 | 0.013 |
| calbindin 2 | *Calb2* | -0.48 | 0.002 | 0.016 |
| hypocretin (orexin) receptor 1 | *Hcrtr1* | 0.47 | 0.0024 | 0.017 |
| glutamate receptor, metabotropic 2 | *Grm2* | -0.46 | 0.0029 | 0.019 |
| 5-hydroxytryptamine (serotonin) receptor 5B | *Htr5b* | -0.46 | 0.0032 | 0.019 |
| glutamate receptor, metabotropic 5 | *Grm5* | -0.46 | 0.0034 | 0.02 |
| purinergic receptor P2Y, G-protein coupled 12 | *P2ry12* | -0.45 | 0.0042 | 0.02 |
| 5-hydroxytryptamine (serotonin) receptor 1A | *Htr1a* | -0.44 | 0.0045 | 0.02 |
| glutamate receptor, ionotropic, kainate 4 | *Grik4* | -0.44 | 0.0045 | 0.02 |
| purinergic receptor P2Y, G-protein coupled, 14 | *P2ry14* | -0.44 | 0.0045 | 0.02 |
| cannabinoid receptor 1 (brain) | *Cnr1* | -0.44 | 0.0051 | 0.022 |
| myelin-associated oligodendrocytic basic protein | *Mobp* | 0.43 | 0.006 | 0.024 |
| glutamate receptor, ionotropic, NMDA2D (epsilon 4) | *Grin2d* | -0.41 | 0.0086 | 0.034 |
| oxytocin receptor | *Oxtr* | -0.40 | 0.01 | 0.038 |
| purinergic receptor P2Y, G-protein coupled 2 | *P2ry2* | -0.39 | 0.012 | 0.044 |
| cannabinoid receptor 2 (macrophage) | *Cnr2* | -0.38 | 0.015 | 0.051 |
| 5-hydroxytryptamine (serotonin) receptor 2B | *Htr2b* | -0.38 | 0.015 | 0.051 |
| opioid receptor, mu 1 | *Oprm1* | -0.38 | 0.017 | 0.054 |
| neurotensin receptor 1 | *Ntsr1* | -0.37 | 0.022 | 0.068 |
| glutamate receptor, metabotropic 1 | *Grm1* | -0.35 | 0.026 | 0.079 |
| pyrimidinergic receptor P2Y, G-protein coupled, 6 | *P2ry6* | 0.35 | 0.027 | 0.079 |
| glutamate receptor, ionotropic, NMDA3B | *Grin3b* | -0.34 | 0.032 | 0.087 |
| glutamate receptor, ionotropic, AMPA1 (alpha 1) | *Gria1* | -0.34 | 0.033 | 0.087 |
| myelin basic protein | *Mbp* | 0.34 | 0.033 | 0.087 |
| glutamate receptor, ionotropic, NMDA2C (epsilon 3) | *Grin2c* | -0.34 | 0.034 | 0.087 |
| 5-hydroxytryptamine (serotonin) receptor 1B | *Htr1b* | -0.33 | 0.037 | 0.089 |
| purinergic receptor P2X, ligand-gated ion channel, 1 | *P2rx1* | -0.33 | 0.038 | 0.089 |
| 5-hydroxytryptamine (serotonin) receptor 3A | *Htr3a* | -0.33 | 0.038 | 0.089 |
| glutamate receptor, ionotropic, NMDA1 (zeta 1) | *Grin1* | 0.32 | 0.047 | 0.11 |
| opioid receptor-like 1 | *Oprl1* | -0.30 | 0.059 | 0.13 |
| glutamate receptor, ionotropic, kainate 3 | *Grik3* | -0.30 | 0.062 | 0.13 |
| histamine receptor H1 | *Hrh1* | -0.30 | 0.062 | 0.13 |
| cholinergic receptor, muscarinic 2, cardiac | *Chrm2* | 0.29 | 0.074 | 0.15 |
| glutamate receptor, metabotropic 4 | *Grm4* | 0.29 | 0.074 | 0.15 |
| cholinergic receptor, muscarinic 1, CNS | *Chrm1* | -0.27 | 0.088 | 0.17 |
| gamma-aminobutyric acid (GABA) B receptor, 2 | *Gabbr2* | 0.27 | 0.093 | 0.17 |
| pleckstrin homology domain containing, family B (evectins) member 1 | *Plekhb1* | 0.27 | 0.092 | 0.17 |
| histamine receptor H3 | *Hrh3* | -0.26 | 0.11 | 0.2 |
| tachykinin receptor 3 | *Tacr3* | -0.25 | 0.12 | 0.21 |
| 5 hydroxytryptamine (serotonin) receptor 4 | *Htr4* | 0.24 | 0.13 | 0.23 |
| opioid receptor, kappa 1 | *Oprk1* | -0.24 | 0.13 | 0.23 |
| vasoactive intestinal polypeptide | *Vip* | -0.22 | 0.17 | 0.28 |
| 5-hydroxytryptamine (serotonin) receptor 3B | *Htr3b* | 0.22 | 0.17 | 0.29 |
| glutamate receptor, metabotropic 3 | *Grm3* | 0.20 | 0.22 | 0.35 |
| dopamine receptor D1 | *Drd1* | -0.20 | 0.22 | 0.35 |
| glutamate receptor, ionotropic, NMDA2A (epsilon 1) | *Grin2a* | 0.19 | 0.23 | 0.35 |
| glutamate receptor, ionotropic, NMDA2B (epsilon 2) | *Grin2b* | 0.19 | 0.23 | 0.35 |

| | | | | |
|---|---|---|---|---|
| dopamine receptor D4 | *Drd4* | -0.19 | 0.25 | 0.38 |
| adrenergic receptor, alpha 2b | *Adra2b* | -0.17 | 0.28 | 0.42 |
| cholinergic receptor, muscarinic 3, cardiac | *Chrm3* | 0.16 | 0.34 | 0.48 |
| glutamate receptor, ionotropic, AMPA2 (alpha 2) | *Gria2* | 0.15 | 0.34 | 0.48 |
| tachykinin receptor 1 | *Tacr1* | -0.15 | 0.34 | 0.48 |
| neuropeptide Y receptor Y1 | *Npy1r* | 0.13 | 0.41 | 0.56 |
| glutamate receptor, metabotropic 8 | *Grm8* | -0.13 | 0.41 | 0.56 |
| melanocortin 3 receptor | *Mc3r* | -0.12 | 0.46 | 0.61 |
| adenosine A2a receptor | *Adora2a* | -0.12 | 0.47 | 0.61 |
| adrenergic receptor, alpha 2a | *Adra2a* | 0.12 | 0.47 | 0.61 |
| cholinergic receptor, muscarinic 4 | *Chrm4* | 0.11 | 0.48 | 0.61 |
| somatostatin receptor 4 | *Sstr4* | 0.11 | 0.49 | 0.62 |
| somatostatin | *Sst* | -0.10 | 0.52 | 0.65 |
| somatostatin receptor 2 | *Sstr2* | -0.10 | 0.53 | 0.65 |
| histamine receptor H2 | *Hrh2* | 0.10 | 0.54 | 0.66 |
| adrenergic receptor, alpha 1d | *Adra1d* | 0.09 | 0.56 | 0.67 |
| glutamate receptor, ionotropic, AMPA4 (alpha 4) | *Gria4* | 0.09 | 0.58 | 0.68 |
| adrenergic receptor, alpha 1a | *Adra1a* | 0.07 | 0.67 | 0.78 |
| arginine vasopressin receptor 1A | *Avpr1a* | 0.06 | 0.7 | 0.79 |
| galanin receptor 1 | *Galr1* | 0.06 | 0.71 | 0.79 |
| opioid receptor, delta 1 | *Oprd1* | 0.06 | 0.71 | 0.79 |
| arginine vasopressin receptor 1B | *Avpr1b* | 0.05 | 0.74 | 0.81 |
| glutamate receptor, ionotropic, AMPA3 (alpha 3) | *Gria3* | -0.05 | 0.78 | 0.85 |
| glutamate receptor, ionotropic, kainate 5 (gamma 2) | *Grik5* | -0.03 | 0.86 | 0.92 |
| dopamine receptor D2 | *Drd2* | 0.02 | 0.89 | 0.94 |
| dopamine receptor D3 | *Drd3* | 0.02 | 0.9 | 0.94 |
| vasoactive intestinal peptide receptor 2 | *Vipr2* | -0.02 | 0.92 | 0.94 |
| calbindin 1 | *Calb1* | -0.02 | 0.92 | 0.94 |
| melanin-concentrating hormone receptor 1 | *Mchr1* | -0.00 | 0.98 | 0.99 |
| adrenergic receptor, beta 1 | *Adrb1* | -0.00 | 0.99 | 0.99 |

**Table 2.** Correlations of 86 brain-related genes to T1w:T2w across 40 mouse cortical areas. We give the Spearman correlation, $\rho_{\mathrm{T1w:T2w}}$, the $p$-value, and the FDR-corrected $p$-value (across 86 independent comparisons). A horizontal line marks the threshold, $p_{\mathrm{corr}} = 0.05$.

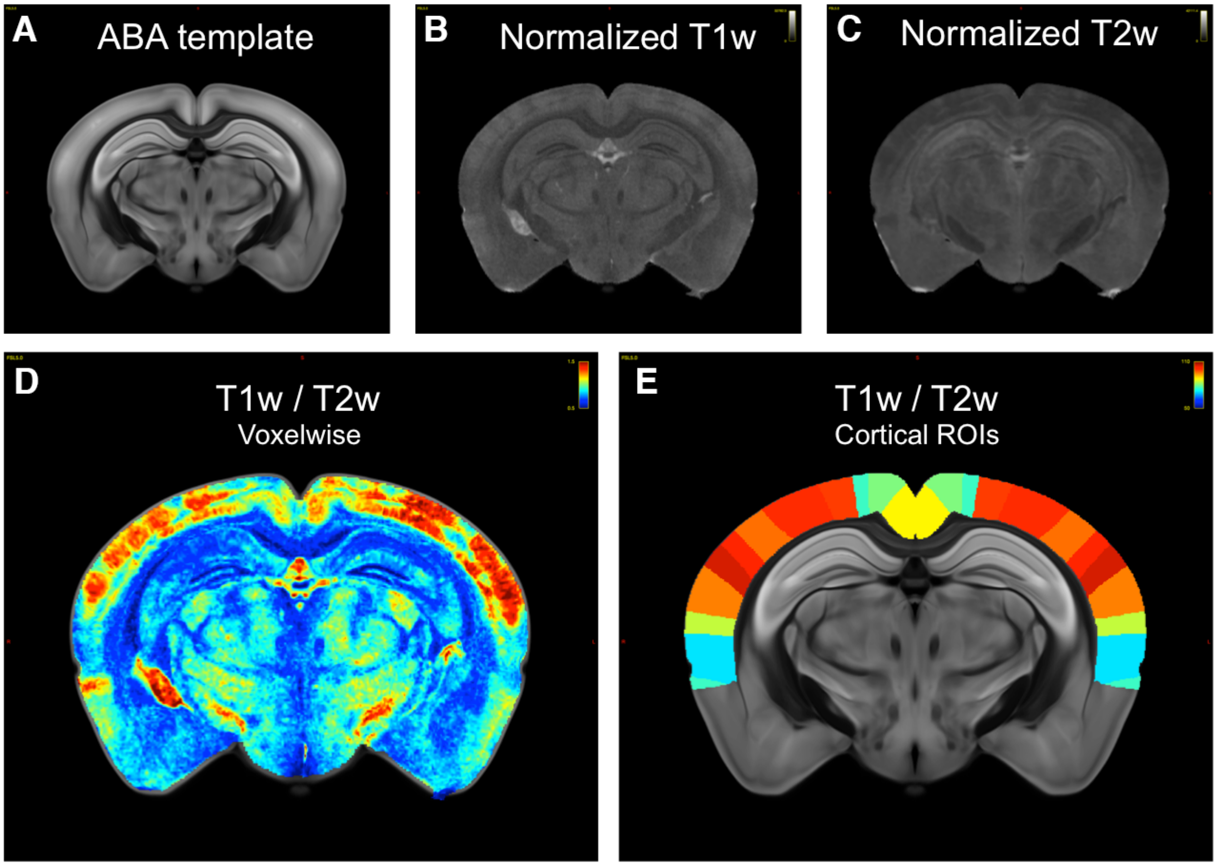


**Fig. 1. Normalization of T1w:T2w between the Allen Mouse Common Coordinate Framework (CCFv3) based on Allen Reference Atlas (ARA) (1) and T1w:T2w maps from Waxholm Space (5). A, B, C** Bias-field corrected T1- and T2-weighted images in Waxholm space were normalized in CCF coordinates via affine and non-linear diffeomorphic deformations. The coronal images, shown at Bregma -3.2 mm, illustrate the quality of the normalization. **D** Voxel-level T1w:T2w images show non-uniform contrasts across the cortex. **E** The Allen Mouse CCF was used to compute T1w:T2w in each cortical region.

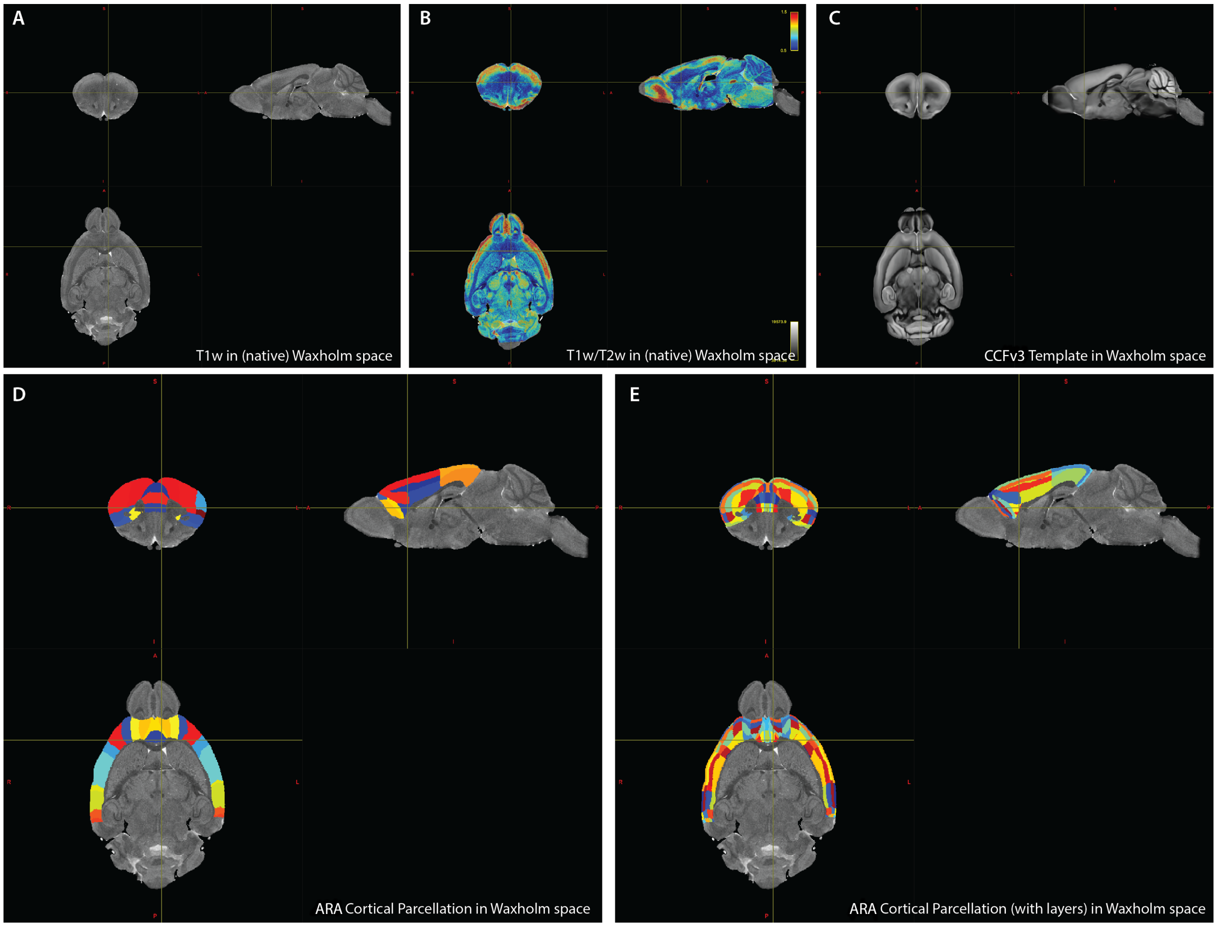


**Fig. 2. Inverted mapping of the Allen CCFv3 and Allen Reference Atlas (ARA) cortical parcellation to Waxholm space (5) demonstrate the accuracy of the normalization**. We plot the measurements and parcellations used here in Waxholm space: **A** T1w (measured natively), **B** T1w:T2w (measured natively), **C** the Allen CCFv3 template (transformed to Waxholm space), **D** the ARA cortical parcellation (without layers) used here (transformed to Waxholm space), and **E** the ARA cortical parcellation (with layers) used here (transformed to Waxholm space). Note that the colors used in **D** and **E** are assigned at random to label discrete cortical parcels.

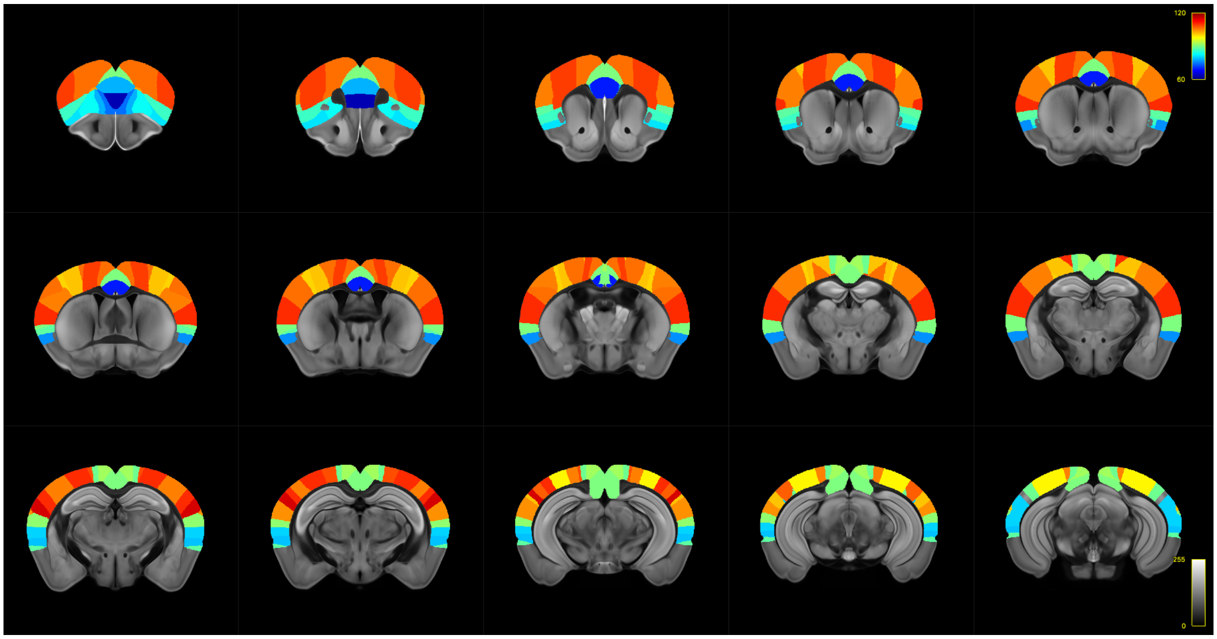


**Fig. 3. T1w:T2w across medial mouse brain sections from the Allen CCFv3** Color indicates the mean T1w:T2w in each of 40 cortical areas, from blue (0.6) through to red (1.2).

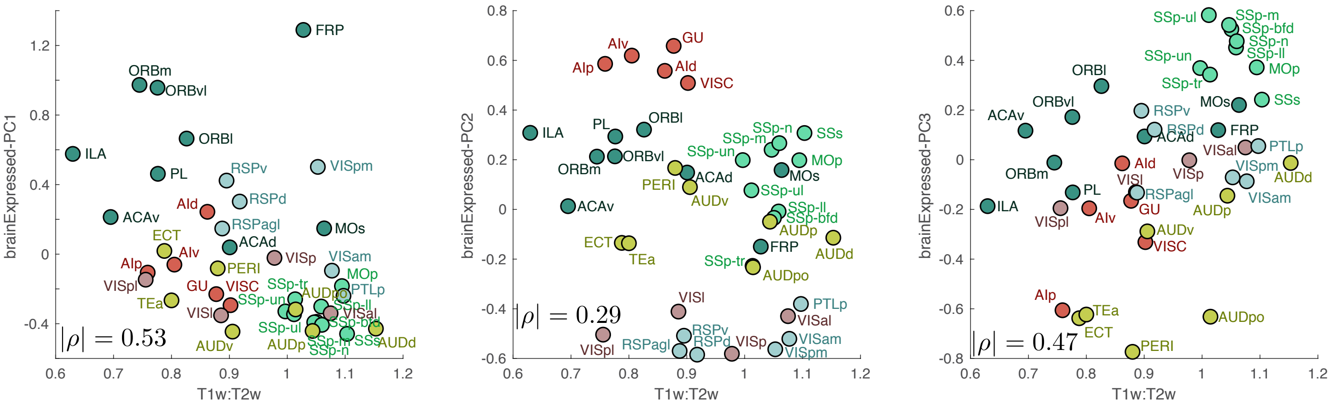

**Fig. 4. Variation of T1w:T2w with the leading three principal components (PCs) of mouse cortical transcriptional maps.** Transcriptional maps were calculated for 1055 brain-expressed genes, which were each normalized using a sigmoidal transformation and then $z$-scored for probabilistic principal components analysis (PPCA). Here we plot the relationship between T1w:T2w (horizontal axis) and each of: **A** PC1, **B** PC2, and **C** PC3. Each PC defines an orthogonal spatial map that incrementally explains maximal remaining variance in the transcriptional maps of all 1055 brain-expressed genes. There is a significant correlation with PC1, $|\rho| = 0.53$ ($p = 6 \times 10^{-4}$) and PC3, $|\rho| = 0.47$, $p = 3 \times 10^{-3}$, but not with PC2, $|\rho| = 0.29$ ($p = 0.07$).

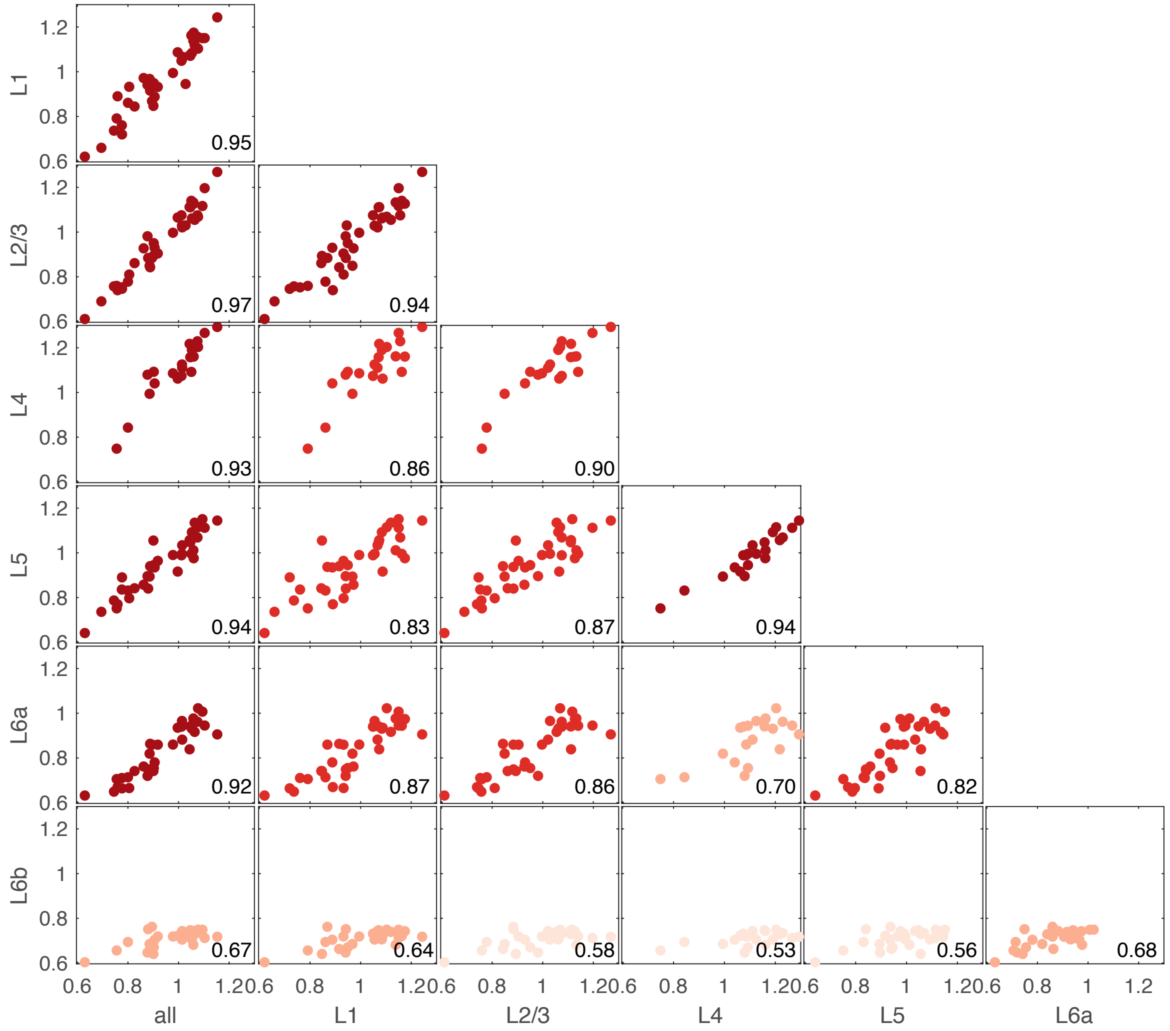

**Fig. 5. T1w:T2w values calculated in each of cortical layers 1–5, are highly correlated to each other and to T1w:T2w values averaged across all cortical layers.** Each scatter plot is of the T1w:T2w averaged across voxels in a single layer (or across all layers, 'all') versus that in another layer. Pearson correlation coefficients are annotated in the bottom-right corner of each scatter plot.

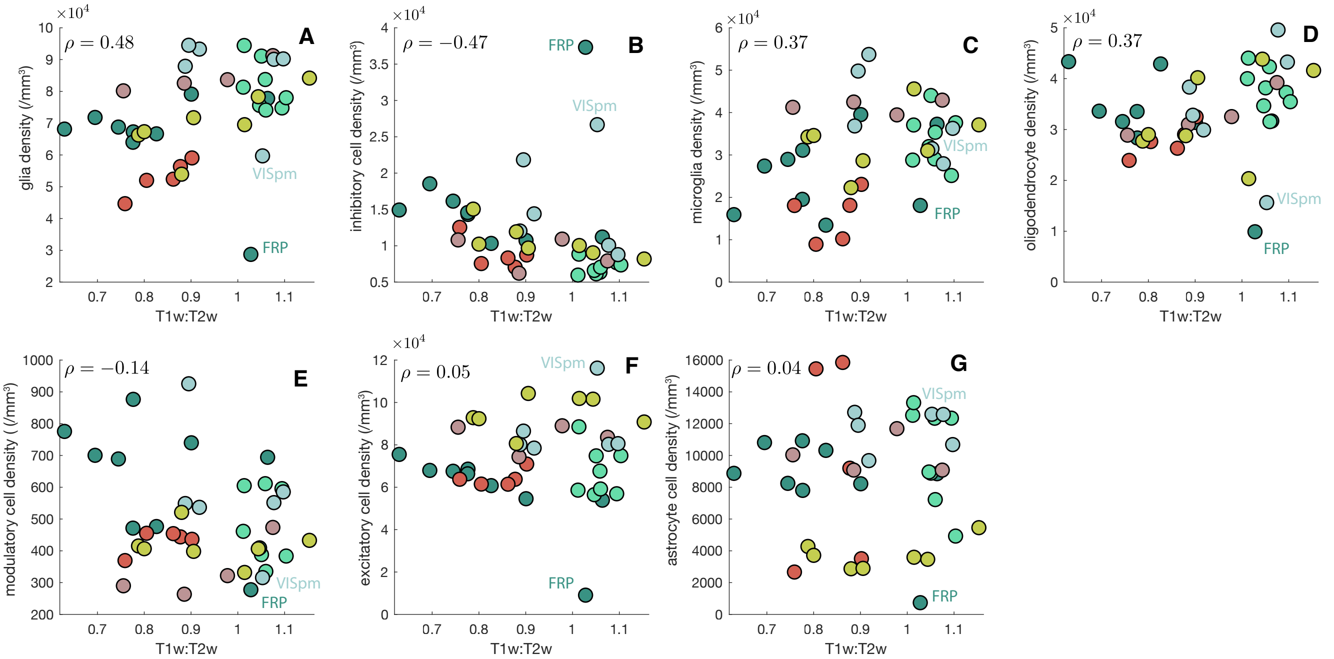


**Fig. 6. Scatter plots of T1w:T2w with cell-density data from (9)** T1w:T2w is plotted against: **A** glia density, $\rho = 0.48$ ($p_{\mathrm{corr}} = 0.01$), **B** inhibitory cell density, $\rho = -0.47$ ($p_{\mathrm{corr}} = 0.01$), **C** microglia density, $\rho = 0.37$ ($p_{\mathrm{corr}} = 0.04$), **D** oligodendrocyte density, $\rho = 0.37$ ($p_{\mathrm{corr}} = 0.04$), **E** modulatory cell density, $\rho = -0.14$ ($p_{\mathrm{corr}} = 0.5$), **F** excitatory cell density, $\rho = 0.05$ ($p_{\mathrm{corr}} = 0.8$), and **G** astrocyte density, $\rho = 0.04$ ($p_{\mathrm{corr}} = 0.8$). FRP (frontal pole) and VISpm (posteromedial visual area), are often outliers to the main trend, and are labeled in each plot.

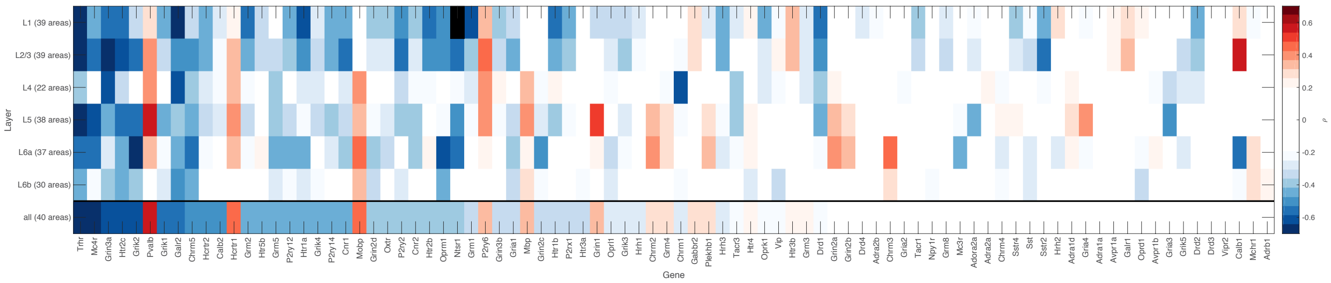


**Fig. 7. Spearman correlations between the expressions of a given gene in a given cortical layer and T1w:T2w across mouse cortical areas** Results are shown for all 86 brain-related genes (missing data shown as black rectangles). One noteworthy pattern is the high positive correlation between T1w:T2w and *Calb1* expression in layer 2/3, and a high negative correlation in layer 6a, but a weak correlation overall; the majority of genes show a similar direction of correlation to T1w:T2w in all cortical layers where there is a correlation.

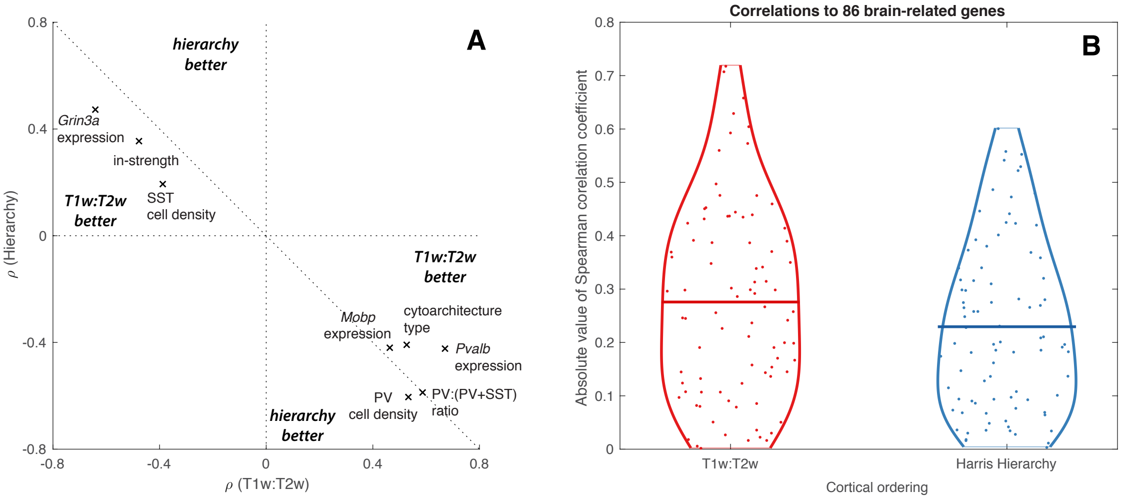


**Fig. 8. T1w:T2w and hierarchy (2) are both markers for diverse cortical gradients, but T1w:T2w shows stronger correlations than hierarchical level in general.** To aid fair comparison between the two measurements, only 35 of our cortical areas with hierarchical level information were used to compute correlations (excluding PTLp, ECT, PERI, AUDv, and GU). **A** Hierarchy and T1w:T2w are negatively correlated to each other, and exhibit broadly similar magnitudes of correlation coefficients, $\rho$, to independently measured structural properties. PV cell density is the only measurement for which hierarchy displays a stronger $|\rho|$ than T1w:T2w. **B** The distribution of correlations to 86 brain-related genes is shown for T1w:T2w and harris hierarchy. The distributions are highly overlapping, but T1w:T2w exhibits a greater average correlation to the transcriptional maps of brain-related genes than hierarchical level (2), which is not significant at the 0.05 level ($p = 0.1$, Wilcoxon rank sum test).

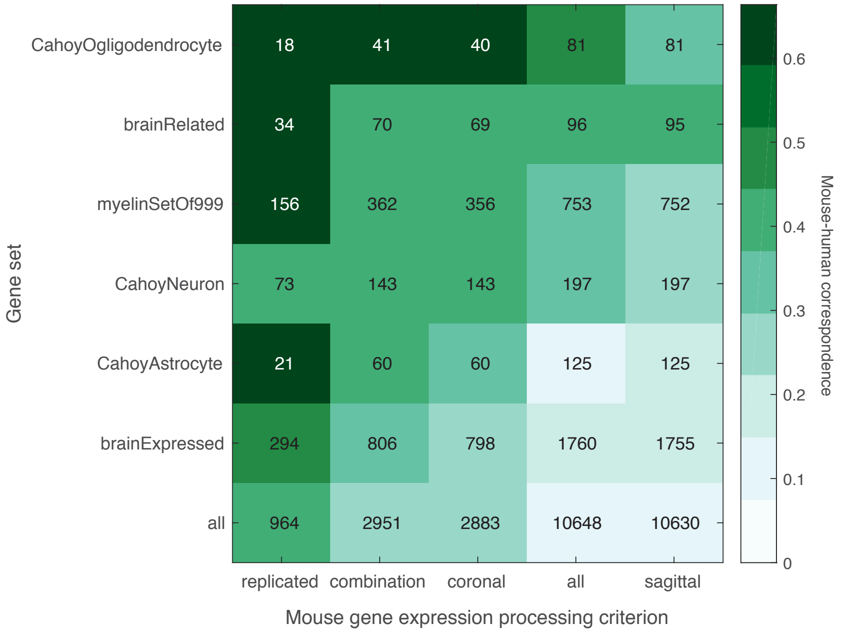


**Fig. 9. Mouse–human consistency in the relationship between gene expression and T1w:T2w increases with data quality and when computed across specific sets of genes.** Rows indicate defined gene sets (defined above); columns indicate the quality-control filtering applied to mouse ISH section data (described above). Rows are ordered by their mean value across columns (decreasing); columns are ordered the same way, by their mean value across rows (decreasing). Color indicates $\tilde{\rho}_{mh}$, the Spearman correlation coefficient computed between $\rho_m$ (correlation between T1w:T2w and transcriptional maps in mouse cortex) and $\rho_h$ (matched human orthologs in human cortex) across the gene set and expression measure of interest. Text annotations label the number of genes used to compute each correlation. The mouse-human correspondence increases along a progression of increasingly high quality criteria for AMBA section dataset inclusion (columns), with the lowest correspondence from using just sagittal section data, and the highest correspondence when we include only genes with multiple experiments with consistent cortical expression patterns. The strongest average mouse-human correspondence is seen for ogligodendrocyte-enriched genes (19), brain-related genes, and myelin-enriched genes, consistent with the known sensitivity of T1w:T2w to myelin (35).
